# MUM, a maternal unknown message, inhibits early establishment of the medio-lateral axis in the embryo of the kelp *Saccharina latissima*

**DOI:** 10.1101/2024.01.07.574535

**Authors:** Samuel Boscq, Bernard Billoud, Ioannis Theodorou, Tanweer Joemmanbaks, Bénédicte Charrier

## Abstract

Brown algae are multicellular photosynthetic organisms that have evolved independently of plants and other algae. Apart from studies on the polarisation of the *Fucus* zygote in the 1990s, little is known about the mechanisms controlling the embryogenesis of these organisms. Here, we studied the determinism of embryogenesis in the kelp *Saccharina latissima,* focusing on the formation of its body axes. This alga initially develops an oblong embryo formed of a cell monolayer, which subsequently thickens; growth axes are then established in temporally distinct stages, starting with the formation of a dominant apico-basal axis. Our experiments focused on the role of the stalk, an empty cell that retains the embryo attached to the maternal tissue, in the development of the growth axes in mature embryos. In response to the removal of the stalk, the embryos developed as a monolayered disc rather than an elongated blade, demonstrating that attachment to the stalk inhibits the formation of the medio-lateral axis at the onset of embryogenesis. In addition, compared with embryos attached to the stalk, the cells of detached embryos were smaller and displayed an altered shape. The topology of the tissue was also disrupted, as cells had a higher number of cell neighbours. Observation of cell division patterns just after removal of the stalk showed that the stalk represses longitudinal cell divisions, thereby reinforcing the establishment of the main apico-basal axis. This unique quantitative study of brown algal embryogenesis revealed that, in kelps, a signal from maternal tissue (MUM for maternal unknown message) is necessary for the establishment of growth axes at the onset of embryogenesis and of the organisation of growing embryonic tissues. In addition, we discovered that, although the stalk persists for several weeks until the embryo reaches at least 500 cells, MUM is emitted in the first 4 days after fertilisation only, before the embryo reaches the 8-cell stage. Finally, transplantation experiments indicated that MUM does not diffuse in seawater, but requires contact between the embryo and the stalk. The potential chemical or mechanical nature of MUM is discussed.

## Introduction

In all organisms, the establishment of spatial axes during embryonic development serves as the fundamental basis for organising the developing body. Despite the immense morphological diversity exhibited by eukaryotes, the majority of organisms can still be characterised by three primary body axes (Anlas and Trivedi, 2021). These axes can be defined based on morphological features (Barthélémy and Caraglio, 2007; Deline et al., 2018; Martinez et al., 2016), or by the presence of molecular gradients within the embryo (Friml et al., 2003; Simsek and Özbudak, 2022). Symmetry-breaking events are crucial for establishing axes within initially homogeneous states and this process is the key behind the facilitation of cell differentiation. However, although polarity is often associated with axis establishment, axial determination can occur independently of polarity in certain symmetrical systems (Cove, 2000).

Brown algae (also named Phaeophyceae, have evolved diverse morphologies (Bogaert et al., 2013; Bringloe et al., 2020; Charrier et al., 2012), ranging from small filamentous forms (e.g. order Ectocarpales) to large multilayered parenchymatous bodies that can reach up to 40 m (e.g. order Laminariales) (Cribb, 1954). They diverged from the ancestors of other extant multicellular organisms at the root of the eukaryotic tree at least 1 billion years ago and emerged as complex multicellular organisms relatively recently, around 250 million years ago (Burki et al., 2020; Kawai et al., 2015). Being evolutionarily distinct from animals, fungi, plants and other algae, brown algae possess unique biological mechanisms (Charrier et al., 2019; Nagasato et al., 2022; Terauchi et al., 2015). They have evolved diverse complex embryonic patterns. For example, contrary to some land plants and metazoans, which develop inside the maternal tissue, brown algae embryos are usually free-growing. Furthermore, after the initial stages of development, they develop simple morphologies. These two features make them easier to access for imaging and manipulation and consequently, brown algae such as *Fucus, Ectocarpus* and *Dictyota* have become developmental models to study polarised cell growth (Bogaert et al., 2017; Goodner and Quatrano, 1993; Rabillé et al., 2019). Reports on *Fucus* and *Dictyota* have shown that the elongation of oospheres occurs only after fertilisation by sperm, but environmental cues such as light can change symmetry, shifting the orientation of the cell before the selection of the axis (Bogaert et al., 2015; Jaffe, 1968).

Unlike these species, no environmental cue such as light polarisation has been reported to affect the formation of the body plans in Laminariales. These algae, like those of orders Desmarestiales and Sporochnales, undergo embryogenesis while being physically connected to their maternal tissue (Fig. 1) (Fritsch, 1945; Klochkova et al., 2019; Wiencke and Clayton, 1990). Therefore, they are powerful models for studying the influence of maternal factors on embryogenesis in brown algae. Although detailed studies have shed light on the impact of maternal tissues on the development of animal and land plant embryos, there is limited documentation for brown algae. During the initial stages of development, the egg cell of *Saccharina,* a member of the Laminariales group, possesses two anchored flagella that facilitate attachment to the gametangia that has differentiated from the filamentous maternal gametophyte (Fig. 1A) (Klochkova et al., 2019). After karyogamy, these flagella degrade (Fig. 1B), but the zygote firmly anchor its base to the female gametophyte, pontentially through the accumulation of glycoconjugates (Klochkova et al., 2019). In 1936, Kanda (Kanda, 1936) was the first to report the abnormal shapes of embryos that happened to detach from the maternal gametophyte.

**Figure 1:**
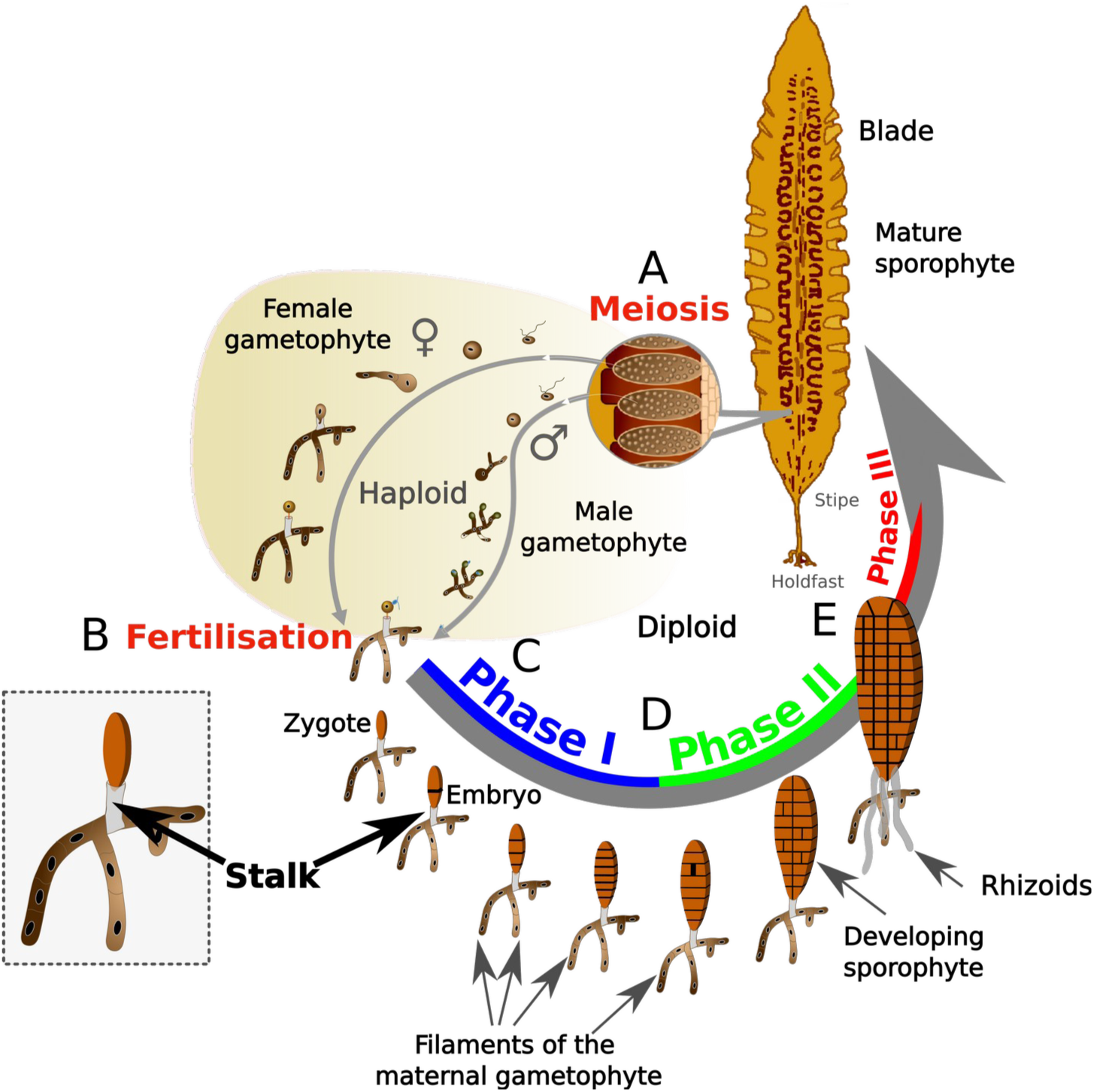
Life cycle of *Saccharina latissima*. (A) In mature sporophyte blades, meiosis occurs inside sporangia and motile meiospores are released in large amounts. The spores fall on the seafloor (or plastic or glassware in the laboratory) and germinate. (B) Spores develop into female and male filamentous gametophytes. When the necessary environmental conditions are met, both types of gametophytes mature and the female gametangium (named oogonium) releases one egg and the male gametangium (named antheridium) releases one sperm cell. (C) After fertilisation, the embryo elongates along the X-axis and initially develops attached to the female gametophyte undergoing a succession of transverse, parallel divisions perpendicular to the zygote axis, up to the formation of a linear stack of 8 cells (Phase I in blue). (D) The embryo starts dividing longitudinally, and continues growth by alternating longitudinal and transverse cell divisions (Phase II in green). (E) After about 15 days after fertilisation, the embryo initiates the first divisions in the Z-axis and undergoes cell differentiation (Phase III in red). The embryo then continues to develop into a mature sporophyte. Colour codes for the three embryogenetic phases are according to Theodorou and Charrier (2023). The insert zooming in on the zygote better displays of the stalk

Recently, the early developmental pattern of the *Saccharina latissima* sporophyte was described in three main steps, corresponding to the establishment of three distinct body plans (Theodorou and Charrier, 2023) (Fig. 1C-E). After fertilisation, the zygote elongates, followed by parallel transverse divisions that lead to the development of an 8-cell stack. Both processes — zygote elongation and transverse divisions — establish and maintain the apico-basal axis, respectively (Phase I, Fig. 1C). Subsequently, the embryo undergoes two-dimensional growth, forming the medio-lateral axis, and growth in these two axes, the apico-basal axis on the one hand and the medio-lateral axis on the other hand forms a small cellular monolayered lamina (also named blade) (Phase II, Fig. 1D). The transition to three-dimensional growth and cellular tissue differentiation (Phase III, Fig. 1E) begins once the blade reaches around 800–1000 cells (Theodorou and Charrier, 2023). In this article, we present a detailed analysis of how the maternal gametophyte controls the establishment of the medio-lateral axis in the early development of the *S. latissima* sporophyte. Using microdissection to separate the maternal gametophyte from the sporophytic embryo, we monitored the development of the early embryos over time. Image segmentation followed by quantitative analyses of the morphological traits made it possible to assess the role of the maternal tissue in the control of body plan formation in the very early stages of embryogenesis.

## Results

To comprehensively investigate the role of maternal tissue in the growth of the embryo, we mechanically detached embryos from the stalk of the female gametophyte at different developmental stages and monitored embryogenesis for up to 14 days in standard culture conditions. Using a micro-needle, we sectioned the embryo from the maternal tissue in Phase I at the egg (E_0_), zygote (E_1_), 2-cell (E_2_), 4-cell (E_4_), 8-cell (E_8_) stages and in Phase II (PhII, corresponding to > 8-cell embryos) (see Theodorou & Charrier, 2023 for the definition of the embryogenetic phases). The impact of the separation of the egg/zygote/embryo (hereafter named E/Z/E) was observed in Phase II, when embryos grow along longitudinal and lateral directions.

### Detachment from the stalk impairs both growth and development of the embryos

We observed a wide range of resulting morphological alterations. A significant proportion of embryos either immediately ceased developing or perished within five days (less than 20% in E_4_ or earlier embryos, not shown). We excluded these embryos from the following analyses.

The remaining living and growing embryos (i.e. at least 80% of the samples) displayed a range of morphogenetic responses illustrated in Fig. 2. Compared with intact embryos (Fig. 2A), those dissected from the stalk before the first cell division (egg E_0_, Fig. 2B; zygote E_1_, Fig. 2C) displayed the most modified patterns. At the egg and zygotic stages, all the embryos showed either delayed growth or altered morphology. The percentage of morphologically altered embryos was higher when the stalk was removed at the egg stage (>80%; e.g. heart-shape embryo, Fig. 2B) than at the zygote stage (40%; Fig. 2C) (Fig. 2H). In contrast, in E_2_and E_4_ microdissected embryos, a significant percentage displayed a developmental pattern comparable to intact embryos (Fig. 2D, and 2E respectively, and Fig. 2H). In the E_8_ and PhII embryos, the effect of detachment from the stalk was weak, with nearly 80% of the microdissected embryos exhibiting a typical developmental pattern (Fig. 2F and G respectively, and Fig. 2H). These data indicate that the connection between the embryo and the stalk before the 8-cell stage is necessary for normal growth and morphogenesis of the embryo.

**Figure 2:**
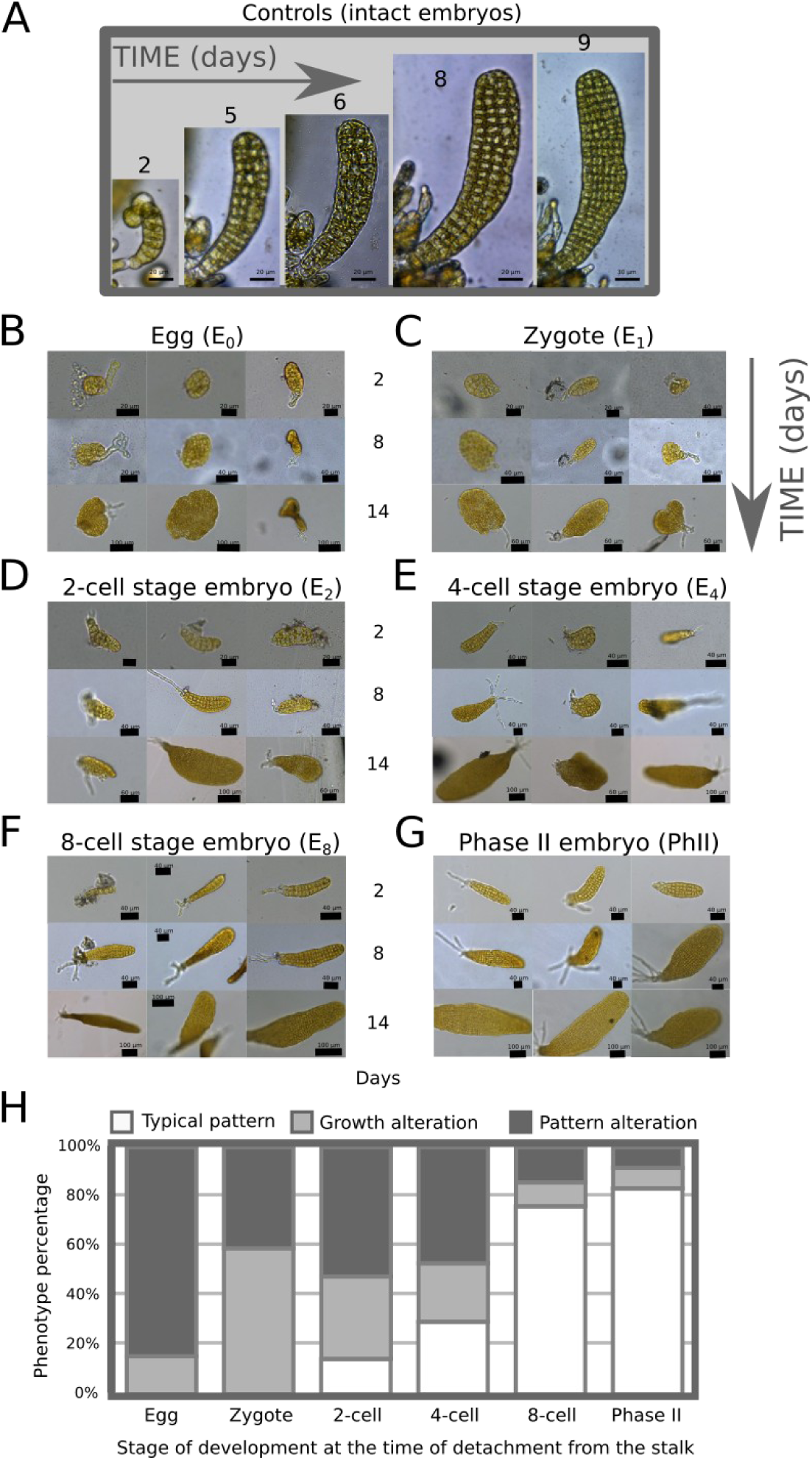
Qualitative impact of embryo detachment from the stalk. Comparison of morphologies in control embryos (A) with growing embryos after being detached from the stalk by microdissection at the egg (E_0_) (B), zygote (E_1_) (C), 2-cell (E_2_) (D), 4-cell (E_4_) (E), 8-cell (E_8_) (F) stage and early Phase II (PhII) (G) and observed 2, 8 and 14 days after microdissection. Three embryos illustrate the representative phenotypes obtained for each microdissection time point. The number of microdissected embryos was n = 4 (A), 100 (B), 100 (C), 80 (D), 80 (E), 40 (F), 40 (G). H) Stacked histogram showing the percentage of each class of phenotype. “Typical pattern” corresponds to morphologies similar to the control; “growth alteration” includes all growth delays and “pattern alteration” corresponds to morphological differences compared with the developmental pattern in intact embryos as displayed in (A). The percentage of typical phenotypes gradually increases with later stage detachment, up to the E_8_stage after which Phase II embryos generally develop similar to intact embryos.

To quantitatively study the impact of dissection from the maternal stalk on the morphogenesis of embryos, we monitored the development of another series of microdissected embryos for 10 days using bright-field microscopy. First, images were captured every day for 4 days, then every two days, and segmented manually (Suppl. Fig. 1). From the segmentation images, quantitative values of several morphometric parameters were obtained from the analysis of the cell outlines using in-house software (Suppl. Table 1) and then compared with statistical tests (see Methods for details; Suppl. Table 2 for the statistical results).

### Detachment from the maternal stalk leads to a disruption of the body plan and reduced growth

To observe how the shape of the embryo is affected by separation from the maternal tissue, we measured the length and the width of PhII embryos containing between 48 and 103 cells ([48:103]). This developmental interval, expressed in the number of cells and not in the number of days of growth, helped to mitigate the effect of the delay in growth due to the stress of the microdissection experiment. Furthermore, neither egg release nor fertilisation are synchronous in *Saccharina*; using embryo age as a reference was therefore impossible with the current protocol of embryo production (Theodorou et al., 2021).

We observed that, in contrast to the intact embryos that display a length/width (l/w) ratio of ∼ 3.5 reflecting their elongated shape, detaching the egg from the maternal tissue resulted in embryos with a disc-like shape (l/w ratio ∼ 1) (Fig. 3A). Further, detachment at different stages between the egg and PhII embryos revealed a gradient in the response: the earlier the separation, the more disc-like the embryo. This morphological response is due to the concomitant modification of two morphological parameters: a reduction in growth along the longitudinal axis (Fig. 3B) (1.9 times less in E_0_ compared with intact embryos; P value < 5.10^-2^) and an increase in growth along the medio-lateral axis (Fig. 3C) (1.3 times more in E_0_ compared with intact embryos; not statistically confirmed). These changes in the direction of growth were accompanied by a significant reduction in surface area of the embryo up to 30% less than the intact embryos (E _0_ and E_1_; P value < 5.10^-2;^ Suppl. Tables 1 and 2).

**Figure 3:**
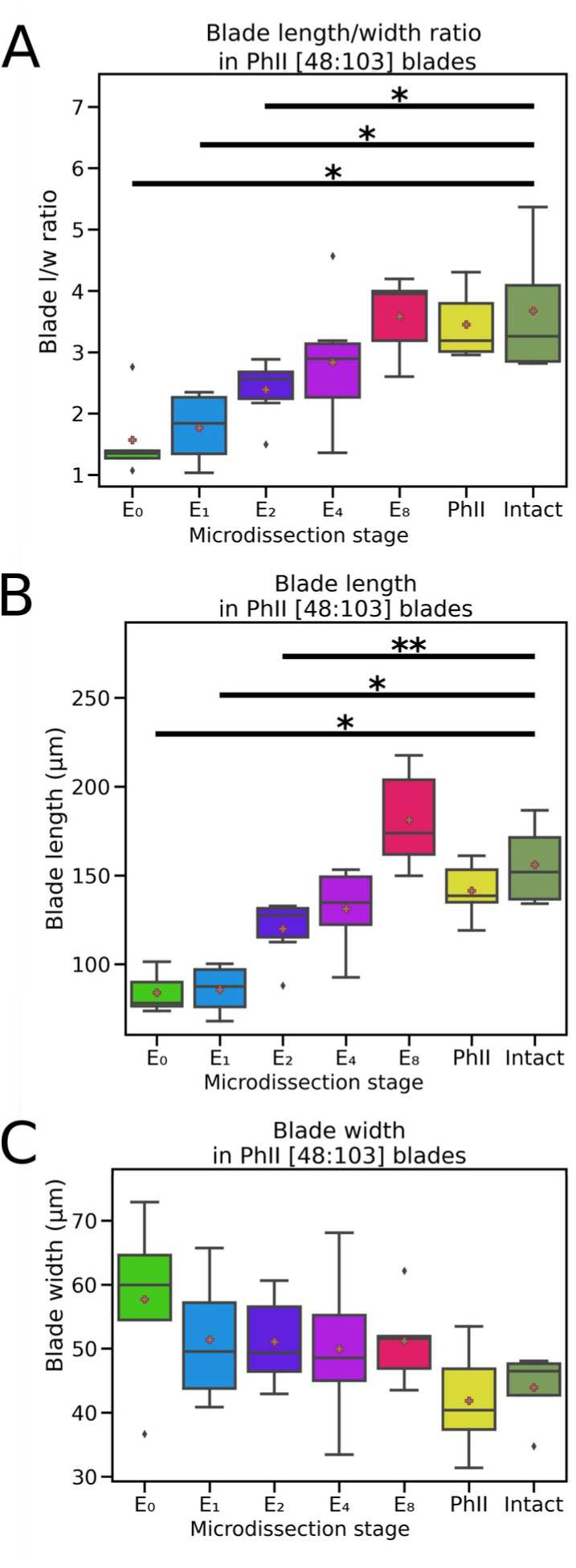
Shape of growing embryos in response to the separation from the maternal tissue. The length and width of Phase II (PhII) embryos made up of 48 to 103 cells [48:103] were measured using in-house blade_painter software and plotted. The middle line is the median, the box includes 25% of the values and the whiskers 75%. Mean is indicated by a plus sign (+). (A) Length/width (l/w) ratio of embryo blade (lamina); (B) Length of the embryo blade: (C) Width of the embryo blade. n=5 for E_0_, 4 for E_1_, 6 for E_2,_ 6 for E_4_, 5 for E_8_, 5 for PhII and 4 for control (Intact). Mann-Whitney test p-value: * <5.10^-2^; ** <1.10^-2^; *** < 10^-3^.

These changes in l/w ratio (and to a lesser extent in surface area) were not only observed in embryos of 43–103 cells, but also in earlier and late embryos (Suppl. Fig. 2; results in late embryos of [104:307] cells were not supported statistically because the sample size was small: 26 embryos vs 34 and 35 in [20:47] and [43:103] cell embryos respectively). This suggests that, as in intact organisms, microdissected embryos maintain the ratio that they had at the time of detachment from the maternal tissue. Thus, the presence of the stalk in the early stages appears to be essential for a long period of embryogenesis and its absence cannot be compensated for until at least the 100-cell stage.

This suggests that a signal related to the maternal stalk controls the development of the embryo. This maternal unknown message (hereafter named “MUM”) operates in real time from the egg stage to the 8-cell stage, albeit decreasing in importance with time, and its impact on embryo morphology lasts up to the 100-cell stage.

### Detachment from the maternal stalk results in altered cell shape and growth

#### Cell growth

We assessed potential alteration of cell growth by measuring cell area using in-house ‘Redraw-blade’ software after segmentation (Suppl. Table 3). Cells of embryos separated from their maternal stalk displayed smaller cells, and the effect was stronger when dissection occurred early on, i.e. at the egg and zygote stages (Fig. 4A). While the average cell area in intact embryos is 69.30 µm^2^, it is 57.51 µm^2^ in E_0_ embryos. Therefore, MUM controls cell size by up to 17% of the reference cell size. This trait was also observed in younger (developmental window: [20–47] cells) and older embryos (developmental window: [104–250] cells) (Suppl. Table 3 and Suppl. Fig. 3). A heatmap of cell area showed that cells with reduced size were not located in specific positions within the embryo, but were spread throughout the embryo (Fig. 4B).

**Figure 4:**
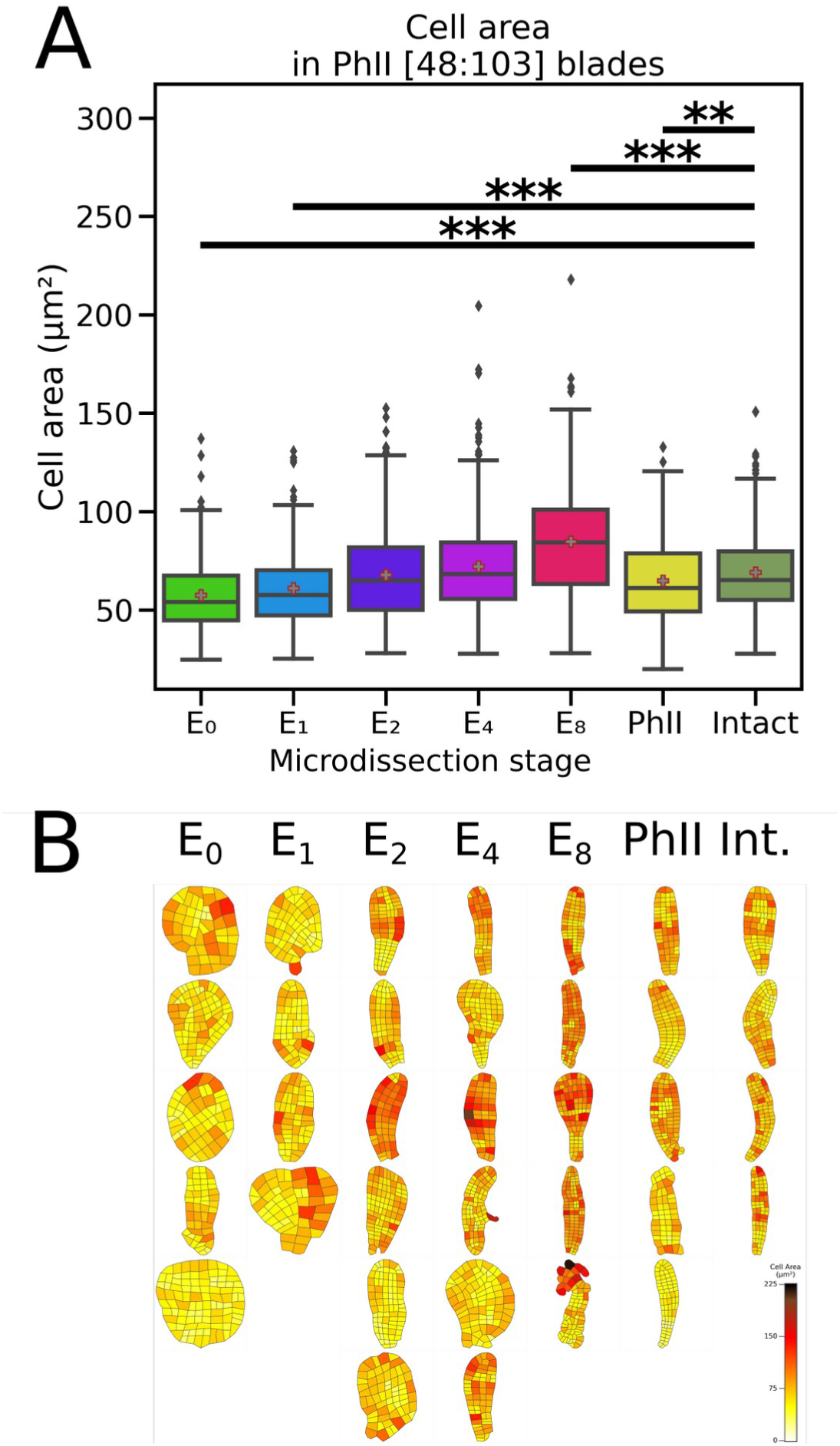
Cell size, expressed as cell surface area of growing embryos detached from maternal tissue. The cell surface areas of 48–103-cell embryos ([48:103]) were measured using in-house blade_painter software and plotted. The middle line is the median, the box includes 25% of the values and the whiskers 75%. (A) Cell area (µm^2^); n=325 for E_0_, 220 for E_1_, 423 for E_2_, 415 for E_4_, 430 for E_8_, 462 for PhII and 306 for control embryos (Intact). *t*-test p-values: * <5.10^-2^; ** <1.10^-2^; *** < 10^-3^. (B) Heatmap of cell area within each embryo considered in (A). Colour scale on the right indicates the range of cell areas observed in the intact and microdissected samples. The size of the image was adjusted so that all blade images have the same dimensions (scales are different).

We addressed whether the decrease in cell size was due to a reduction in cell growth or to a faster cell division rate. From another series of microdissected E/Z/E monitored with images taken every 2 h, we measured the cell division rate and compared it with that of intact embryos (Suppl. Fig. 4). In these microdissected E/Z/E, cell division took place at the same pace as in intact embryos (Fig. 5). As a result, we hypothesise that in the absence of MUM cells are smaller because they expand less while maintaining an unchanged cell division rate. Therefore, MUM appears to promote cell growth independently of cell division.

**Figure 5:**
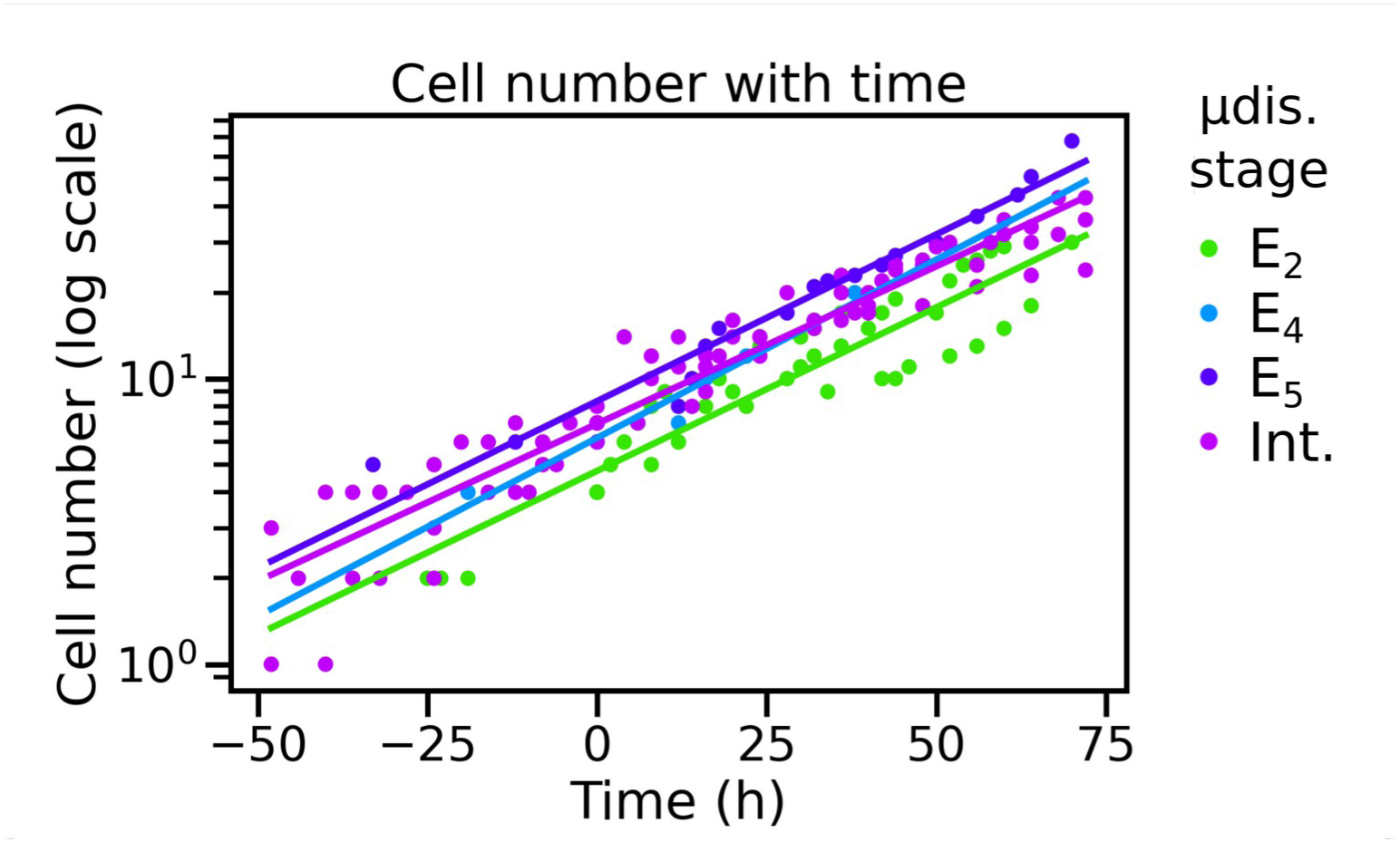
Cell division rate over time in growing embryos detached from maternal tissue. The number of cells of intact (Int.) or microdissected (µdis.) embryos imaged every two hours (as shown in Suppl. Fig. 4) was plotted over time (total duration of the time-lapse: 11 days). The slope of the logarithm of this number of cells represents the growth rate. n=3 for E_2_, 1 for E_4_, 1 for E_5,_ and 5 for Int.

#### Cell shape and impact on neighbouring cells

We defined a rectangularity factor to quantify cell shapes in microdissected and intact embryos. The rectangularity factor was calculated based on a minimum bounding rectangle fitted around the polygonal contour of each cell. The factor equals one when the cell shape, taken in 2D, is a perfect rectangle (all angles joining two sides are 90°), and tends to zero as the angles deviate from this value, corresponding to a “flatter” quadrilateral or a more irregular polygon.

At the 48–103 cell stage (Suppl. Table 3), we noted that cells from microdissected embryos were less rectangular than those of intact embryos (Fig. 6A). The difference was small but significant for all microdissection stages. Similar results were obtained for early and late PhII embryos (Suppl. Fig. 5). In intact embryos, cells with a shape distinct from a perfect rectangle were mainly located in the apex of the blade, where the shape of the blade is curved. In the microdissected embryos, irregular polygons were observed scattered throughout the disc-like blade, with no specific location (Fig. 6B).

**Figure 6:**
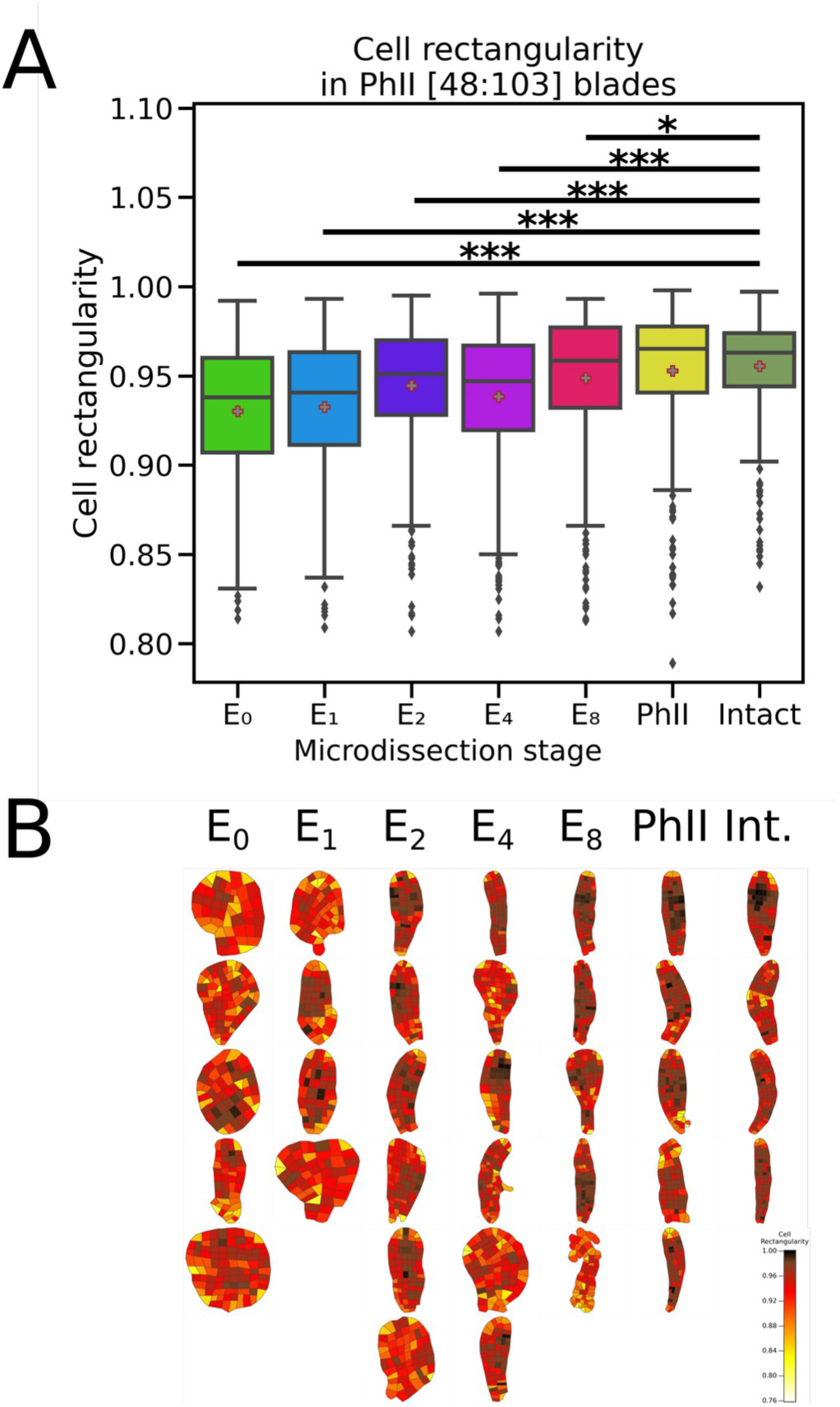
Cell shape in growing embryos detached from maternal tissue. The shape of cells of 48–103-cell embryos ([48;103]) was quantified. (A) The shape of the cell, expressed by a rectangularity factor, which assesses the level of rectangularity of a parallelepiped (cuboid cell seen in 2D). The rectangularity factor is 1 for a parallelepiped with perpendicular sides and < 1 for all other cases. This factor is the lowest for parallelepipeds with very acute angles. n=325 for E_0_, 220 for E_1_, 423 for E_2_, 415 for E_4_, 430 for E_8_, 462 for PhII and 306 for control embryos. *t*-test p-values: * <5.10^-2^; ** <1.10^-2^; *** < 10^-3^. (B) Heatmap of the rectangularity factor. The colour scale on the right indicates the range of values. The size of the image was adjusted so that the image of each blade has the same dimensions as in the figure.

We studied the impact of this change in cell shapes on the topology of the tissue. Specifically, the number of cell neighbours was calculated for each cell. Intact embryos were excluded from this quantification, because they were segmented manually with a different technique that produced more “broken” outlines and resulted in an overestimation of the number of cell neighbours. In PhII embryos, the number of cell neighbours was distributed into two main groups (Fig. 7): cells at the periphery of the blade that were surrounded by three neighbours (one above, one below and one to the side) and cells located inside the blade surrounded by four neighbours (above, below and both left and right sides) (Fig. 7B). In the PhII embryos, only 8.3% of cells had five neighbours, whereas in E_0_ and E_1_ microdissected embryos, 15.7% and 21.4% of cells, respectively, were surrounded by five neighbours (Fig 7A). Similarly, 1.4% of cells of PhII embryos had six cell neighbours, reaching 4.9% and 3.6% in E_0_ and E_1_, respectively. The heatmap shows that these cells were located within the lamina tissue with no specific position related to the apico-basal or medio-lateral axes. Altogether, the general distribution of the number of cell neighbours was significantly different in E_0_, E_1_, E_2_ (P value < 10^-3^; Suppl. Table 2) and E_4_ (P value < 5.10^-2^; Suppl. Table 2) embryos compared with PhII embryos. Similar results were obtained at earlier and later developmental stages (Suppl. Table 3) (Suppl. Fig. 6). Observations on 103–307 cell stage embryos show that this response persists late in Phase II, but only in disc-shaped embryos (Fig. 7B, right, E_4_ microdissection stage). Therefore, removal of the stalk at the 4-cell stage or earlier results in a long-term alteration of the spatial arrangement of cells within the monolayer lamina.

**Figure 7:**
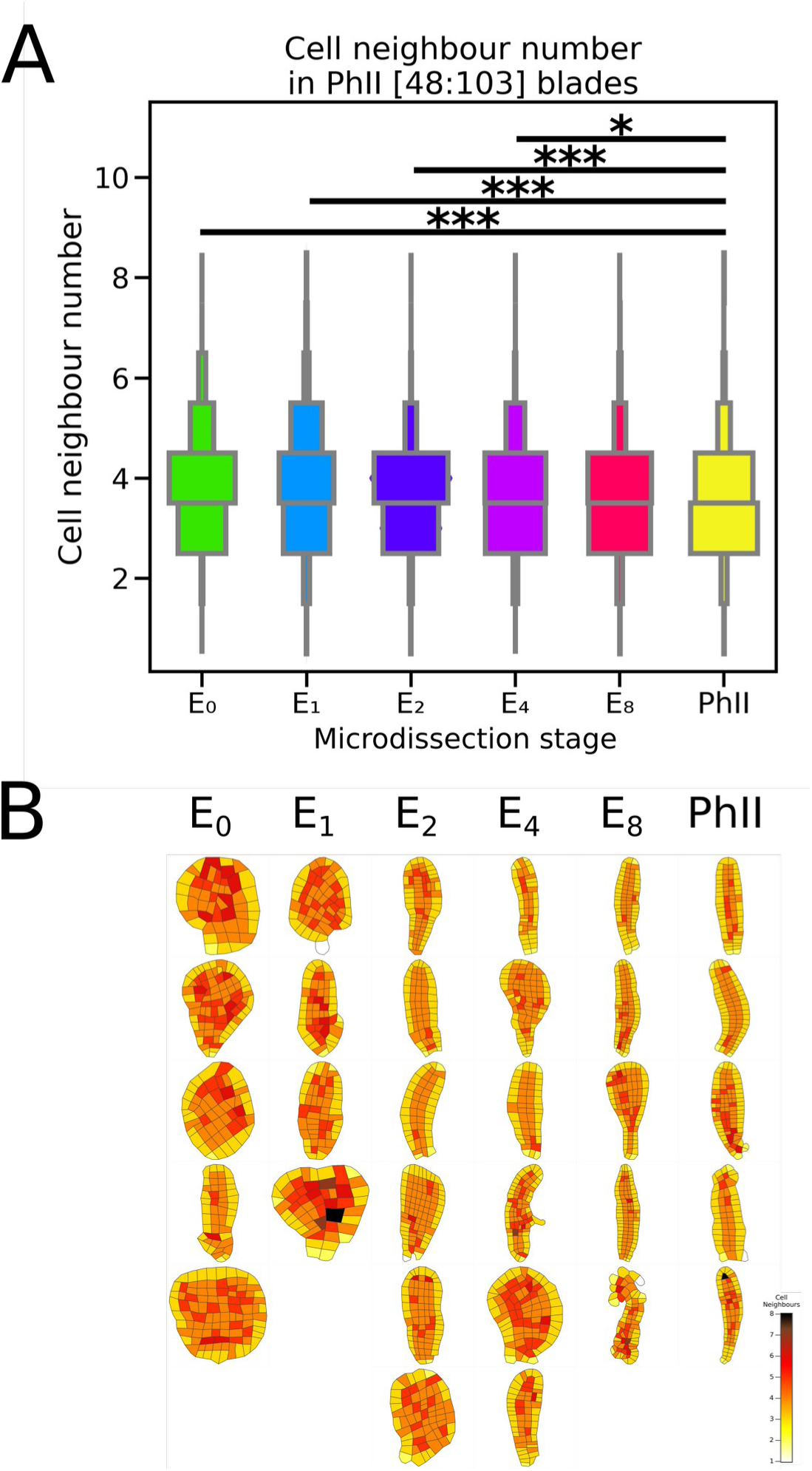
Topology of the embryo blade a in growing embryos detached from maternal tissue. The topology of the growing embryo was studied by counting the number of cells surrounding each cell of 48–103-cell embryos ([48:103]). (A) The number of neighbouring cells was plotted. n=325 for E_0_, 220 for E_1_, 423 for E_2_, 415 for E_4_, 430 for E_8_, 462 for PhII (used as a control). *t*-test p-values: * <5.10^-2^; ** <1.10^-2^; *** < 10^-3^. (B) Heatmap of the number of neighbouring cells for each blade cell. The colour scale on the right indicates the range of values. The size of the image was adjusted so that the image of each blade has the same dimensions as in the figure.

In summary, the blades from which the stalk was removed before the 8-cell stage displayed an altered topology with a higher proportion of cells with more than four neighbours, presumably because these cells were irregularly shaped. In addition, our results showed that the cells were smaller than in intact embryos, which may further contribute to an altered cell arrangement within the blade. Nevertheless, the cells maintain their l/w ratio, at least up to the 100-cell stage (not shown; Suppl. Table 2 for statistical results).

### The disruption of the body plan is due to early longitudinal cell divisions in Phase I

So far, we have shown that the separation of the E/Z/E from the maternal filament resulted in morphological defects in PhII embryos. Namely, the embryos developed as disc-like blades, with smaller, irregularly shaped cells, which are haphazardly arranged within the lamina. To determine whether MUM controls cell shape and size as the main targets, thereby affecting embryo shape, or whether MUM controls embryo shape, which in turn affects cell shape and size, we looked at the earlier steps of embryo development. We observed that the detachment of embryos from the maternal stalk modified the orientation of their initial cell divisions. This effect was stronger when detachment occurred in the E_0_ and E_1_ stages (Fig. 8). Longitudinal cell division parallel to the position of the stalk took place soon after the separation of the egg or zygote from the maternal tissue (Fig. 8B), whereas in embryos microdissected at the 8-cell stage or in intact embryos (Fig. 8A), longitudinal cell division rarely took place before the 8-cell stage. This difference in the timing of longitudinal cell division suggests that MUM controls the orientation of the cell divisions in the very initial stages of embryogenesis. Nevertheless and interestingly, even in E_0_ and E_1_, the first cell division is always transverse.

**Figure 8:**
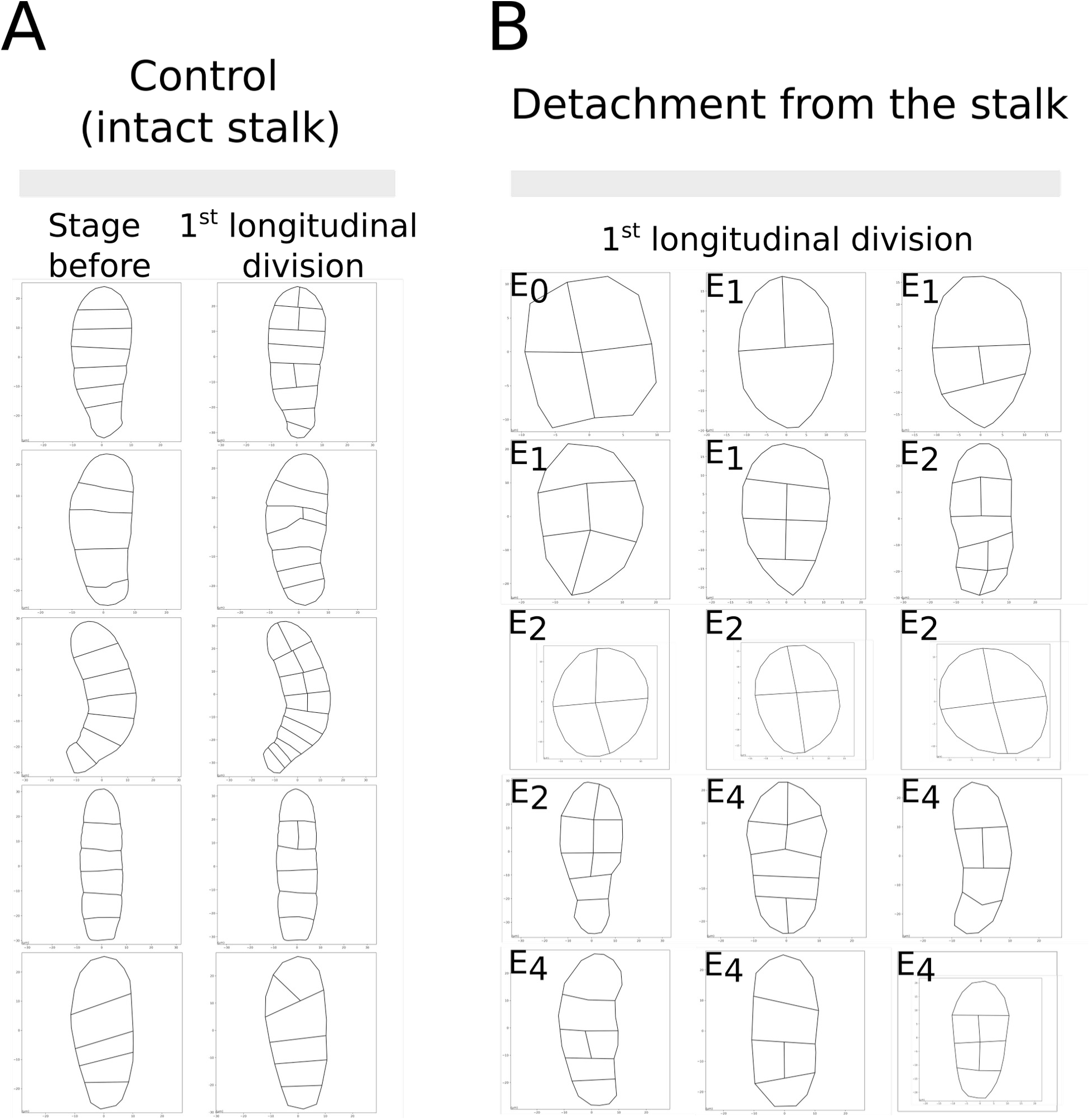
Early longitudinal cell division in growing embryos detached from maternal tissue. The pattern of the embryo blade is displayed at the stage of the occurrence of the first longitudinal cell division(s). (A) Patterns one time-lapse step before (left) and at the emergence (right) of the first longitudinal cell division are displayed for control organisms for which five time-lapse experiments were monitored. (B) Pattern of segmented embryos, which underwent detachment from maternal tissue at the E_0_, E_1_, E_2_ or E_4_ stages (stage of microdissection is indicated in the left-hand corner of each picture). Longitudinal cell division takes place much earlier than the 8-cell stage and often just after stalk removal, except for E_0_ and E_1_, in which the first cell division is always transverse, as in intact embryos. Scales (µm) are indicated in the X and Y axes of each frame.

This result made it possible to identify the cause of the disorganisation of the embryo morphology observed in Phase II. In early Phase I, the primary longitudinal, apico-basal axis is not yet fully established, and the occurrence of longitudinal divisions perpendicular to this axis in the absence of MUM results in embryo growth along the medio-lateral axis. This contrasts with intact embryos exposed to MUM, where a linear stack of eight cells is produced before the second body axis is established.

It is interesting to note that, in the absence of MUM, only divisions parallel or perpendicular to the stalk (prior to dissection), and no oblique orientation, were observed in the growth steps following stalk removal (Fig. 8). Therefore, MUM does not control the orientation of cell division *per se*, but only the conditions of when and where longitudinal divisions occur.

### MUM is a short-range signal

#### What is MUM and how does it act?

##### MUM is a local signal

First, we tested whether MUM is a diffusible factor by immersing detached E/Z/E into Petri dishes containing intact fertile gametophytes. We did not observe any modification in embryo development compared with experiments in which embryos were isolated from the maternal tissue and transferred into a Petri dish with fresh seawater (not shown). This result suggests that there is no molecular or chemical compound excreted from the maternal gametophyte that diffuses in seawater and controls the development of the embryo. Secondly, we investigated whether maternal tissues distant from the embryo, but belonging to the same gametophyte as the embryo, were necessary for proper embryo development. This experiment aimed to test whether a signalling molecule is transported symplastically or apoplastically through the maternal gametophyte filament up to the embryo. We cut or damaged cells of the gametophyte that were directly adjacent to or separated by a few cells from the stalk and we monitored the development of the embryos for the following two weeks. These zygotes developed similarly to control zygotes (Fig. 9). Most zygotes developed a stack of up to eight cells before their first transverse anticlinal division and continued their growth in 2D, forming long ovoid blades without any bleaching, which is a sign of stress. Therefore, the proper development of these embryos suggests that maternal cells close to or adjacent to the embryo do not control its early development.

**Figure 9:**
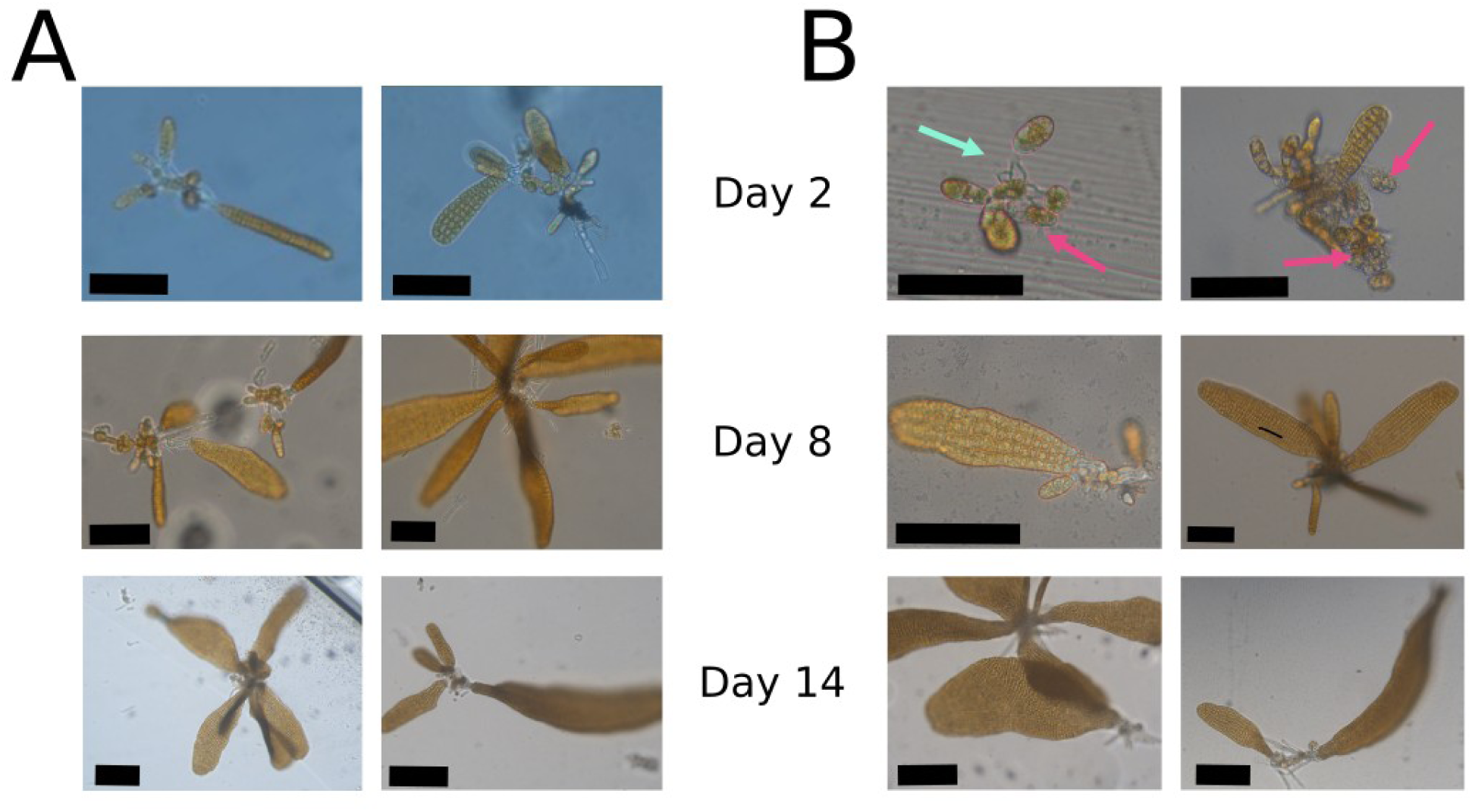
Lack of morphological impact of damaged cells close to the embryo. Because in our experiment, gametophytes are made up of about 10–15 cells, several eggs were produced at different times. Hence, after fertilisation, embryos grew within a “bouquet” of several embryos all attached to a single maternal gametophyte. Embryos were monitored 2, 8 and 14 days after damaging the tissues. (A) Normal conditions of growth. (B) Cells of neighbouring embryos of the same “bouquet” (pink arrows) or of the maternal gametophyte (light blue arrow) were crushed or pierced with the microdissection pipette. Two examples of each condition are shown at each time point. Scale bars are 50 µm for 2 days, 100 µm for 8 days and 200 µm for 14 days post cell damage. n=4 gametophytes bearing several embryos at each time point (different embryos are photographed between time points).

These two experiments support that MUM is neither a signalling factor diffusing in seawater to the E/Z/E, nor a signalling factor transported from cell to cell in the female gametophyte to the stalk. Therefore, MUM is a signalling factor related to the stalk itself.

##### The stalk is a dead structure

The literature indicates that, in Laminariales, the oocyte is released from the mature oogonium through the ejection of the cellular content of the oogonium from an opening at its apex, thereby producing a protoplast (the egg) and a remnant, inert cell wall of the oogonium (Lüning, 1981; Klochkova et al., 2019). We confirmed that the remnant cell wall of the oogonium in *Saccharina latissima* is a dead structure by staining it with Trypan blue (TB), which is a negatively charged dye that living cells with intact membranes do not take up (Farah et al., 2015). TB accumulated in the stalks as well as in dead gametophyte cells, that were used here as controls (Fig. 10). Interestingly, TB staining was stronger at the apex of the stalk near the connection to the embryo. Previous experiments have shown that the stalk interior is highly viscous, and we have observed this viscous material leaking from the stalk when pierced upon mechanical or laser manipulation (Boscq et al., 2022). Therefore, in addition to confirming that the stalk is a dead structure, TB showed that its interior is dense and heterogeneous.

**Figure 10:**
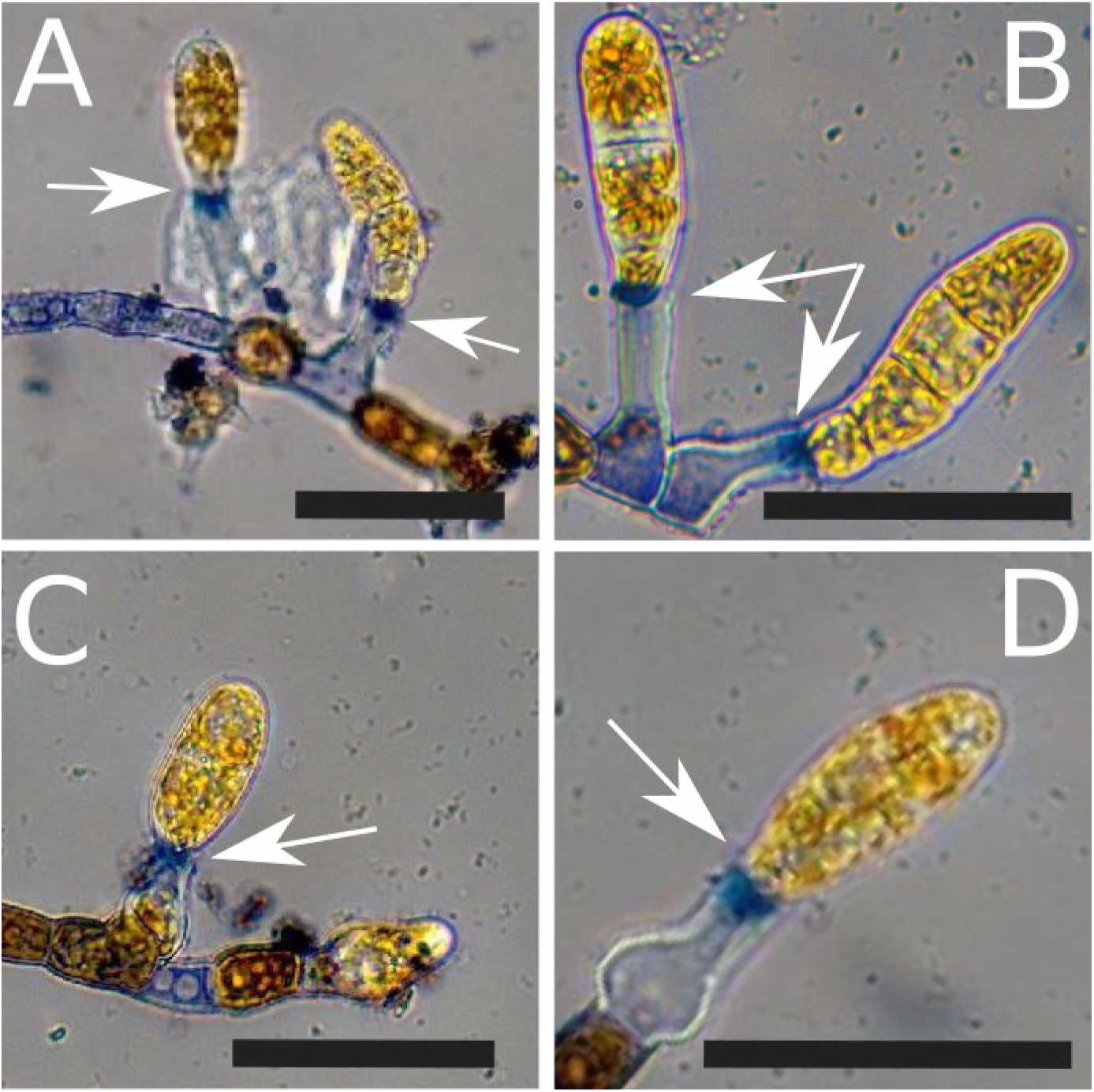
Trypan blue staining of the stalk. Embryos are fixed to the stalk (white arrows). Tissue exposed to Trypan blue showed blue precipitates in the stalk. Staining was stronger at the apex of the stalk than at its base. While some empty cells of the maternal gametophyte were also stained, indicating that they are dead (A, C), live cells remain brown (early stage embryos and gametophyte cells). Four examples are shown (A–D). Scale bar, 50 µm.

##### The stalk is necessary for the control of the formation of the longitudinal axis

It has long been reported that eggs naturally detached from the stalk develop abnormally (Kanda, 1936) (Sauvageau, 1918) (p. 155 on *Laminaria flexicaulis*, former name of *Laminaria digitata*, which also belongs to order Laminariales). Growth is delayed (Fig. 11A) and the distinction of axes is lost. Same results were obtained when flagella were removed (Klochkova et al., 2019). Because these authors did not observe cell divisions at very early stages of embryogenesis, we aimed to repeat the experiments by monitoring cell division just after the egg separation from the stalk. When embryos were forcefully pulled apart, instead of delicately separated from the top part of the stalk by microdissection, we observed that cell division was erratic (Fig. 11B, Suppl. Movie 1). Embryos divided in all spatial directions and presented bulging cell shapes. This response is different from that observed when the stalk is sectioned.

**Figure 11:**
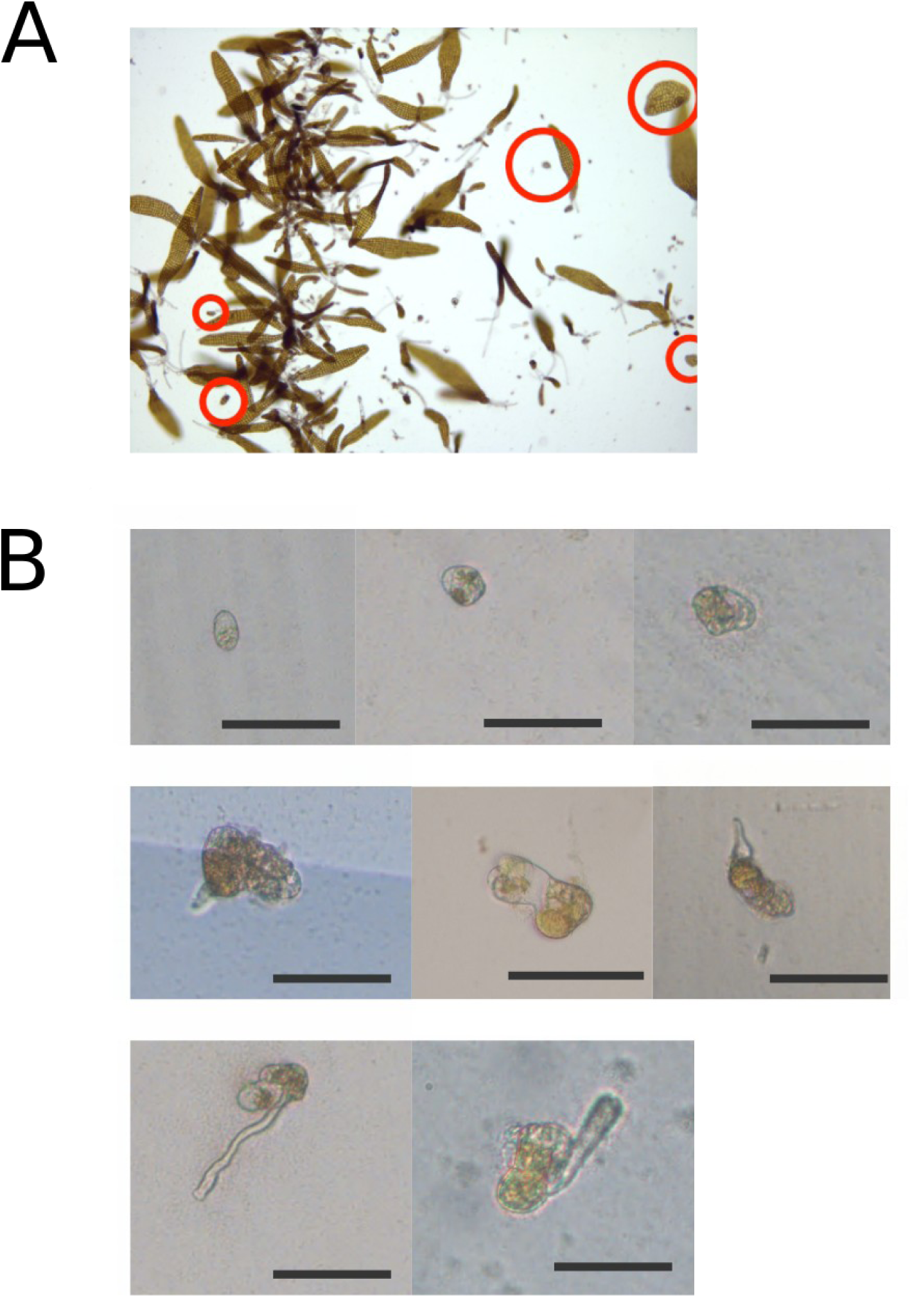
Abnormal embryos growing from eggs naturally detached from the maternal gametophyte. Eggs that were detached from the maternal tissue by sea currents or after shaking the Petri dish in the present case, developed into embryos with major growth delays (A, embryos circled in red) and loss of the main body axes due to erratic orientations of cell divisions (B; 8 embryos are shown). Scale bars, 100 µm.

## Discussion

In this study, we investigated the critical factors that influence the establishment of body axes during embryogenesis in brown algae. Our findings provide insights into the role of the maternal tissue, particularly the stalk, in establishing the two primary axes of the body plan.

The first body axis of the embryo is the longitudinal axis (X-axis; Theodorou and Charrier, 2023), reportedly established upon fertilisation of the egg of Laminariales (Kanda, 1936). After fertilisation, the egg of *Laminaria angustata*, another member of the Laminariales like *Saccharina latissima*, accumulates vacuoles in the future apical region and chloroplasts in the basal region close to the female flagella (Motomura, 1990). However, in rare cases, we observed the release of elongated eggs (not shown), suggesting that egg elongation can occur before fertilisation (also reported in Lüning, 1981). After fertilisation, cells divide transversally first, then longitudinally. Interestingly, different genera in order Laminariales display different distributions of transverse and longitudinal cell division orientations (Kanda, 1936; 1938; 1941; Sauvageau, 1918), resulting in embryos with varied l/w ratios. Therefore, control of Y-axis establishment is a common but plastic trait among individual embryos and among Laminariales species.

### The formation of the medio-lateral axis is controlled by a maternal unknown message (MUM)

We showed that detaching the embryo from the maternal tissue in the very early stages of embryogenesis results in embryogenesis defects. The embryo growth axes were disturbed. Intact embryos grew in length, but embryos separated from their parental tissue tended to grow as a disc, thereby losing their anisotropic shape. Looking at the very first steps of embryogenesis, we noticed that longitudinal cell divisions occurred earlier than in intact embryos. Intact embryos first divided only transversally up to the 8–10-cell stage, whereas the microdissected embryos initiated longitudinal cell division as soon as the 2-cell stage. Noteworthily, in the latter case, longitudinal cell division never occurred before the 2-cell stage, suggesting that the egg axis is aligned with the stalk prior to that stage, perhaps before the oocyte is released. Therefore, while the X-axis of the zygote seems to be established at the egg stage at the latest, its maintenance and the establishment of the Y-axis depends on the contact of the sporophyte with the female gametophyte. As well as controlling the formation of body axes, MUM could be a trophic factor, as microdissected embryos have smaller surface areas than intact embryos.

### MUM is a local signal emitted from the stalk at the very beginning of embryogenesis

For most morphometric and topological parameters observed in this study, the effect of MUM was stronger at the egg and zygote stages than at the 2-cell, 4-cell, 8-cell and early Phase II embryos. Furthermore, the embryo response was gradual from the egg-stage to the 8-cell stage. When the embryo has reached Phase II, detachment from the stalk no longer has any effect. Therefore, MUM acts in the very first step of the embryogenesis.

Furthermore, sectioned embryos growing together (in the same Petri dish) with the female gametophyte they were sectioned from, or with distinct intact female gametophytes, developed in a similar way to embryos detached from the maternal tissue and transferred to a distinct Petri dish. Therefore, MUM does not appear to be a signalling factor able to diffuse in seawater from fertile female gametophytes, but instead requires a direct contact with the E/Z/E. Can MUM diffuse from the filament of the maternal gametophyte? Very surprisingly, in nature, normal embryos can grow from a maternal gametophyte made up of only one cell (Kanda, 1936). Similarly, in the lab, meiospores collected from a wild fertile sporophyte that were immediately exposed to white light germinate and grow into one cell and immediately produce an oogonium (Lüning, 1981; Theodorou et al., 2021). In both cases, the embryos grow in the same manner as those attached to a multicellular gametophyte. Therefore, MUM cannot be a signal diffusing from the filaments of the maternal gametophyte, which suggests that MUM acts locally at the level of the stalk.

Interestingly, in embryos that have grown up to 1000 cells (end of Phase II), the base of the blade remains the narrowest part of the embryo. Observations show that longitudinal division is less frequent in this region (not shown). However, although the stalk physically remains present up to the end of Phase II when the embryo reaches around 1000 cells, the impact of MUM is reduced to the very early stages of Phase I as shown in this study. Therefore, another factor must relay the inhibitory effect of MUM on the widening of the base after the 8-cell stage.

### Is MUM the egg flagella embedded into the stalk?

Female centrioles, through their role in the formation of flagella anchored to the stalk, are thought to be responsible for the formation of the embryo’s longitudinal axis (Klochkova et al., 2019). When the egg is devoid of flagella (e.g. manually pulled away from the stalk, or separated by shaking the flask), most eggs stop growing (Klochkova et al., 2019). Furthermore, it is thought that the phenotypes observed by (Kanda, 1936) and (Sauvageau, 1918) on detached embryos (supposedly at the egg stage), which are to some extent reminiscent to those observed in this study, may be due to a loss of flagella at the egg stage. Hence, MUM would be related to — or even be — the flagella. However, several arguments reject this hypothesis. First, the microdissection method that we used, even if not precise, usually left some of the stalk attached to the E/Z/E, and therefore, the flagella were most likely not sectioned and were, at worse, only damaged. Secondly, the female centrioles, which are necessary for flagella, degrade as early as the zygote stage (Motomura, 1990; Motomura, 1991), but the phenotypes that we report were observed when the stalk was detached up to the 4-cell stage, and even the 8-cell stage (cell rectangularity). Some authors have shown that flagella can remain up to the 2-cell stage (Klochkova et al., 2019). However, it must be a rare case corresponding to the flagella devoid of centrioles trapped in the mucoid component of the stalk. Therefore, the persistence of MUM action throughout most of Phase I embryogenesis excludes that MUM is the flagellar structure itself. Male centrioles replace the female centrioles at the zygote stage (Motomura et al., 2010). In our study, ablation of the stalk was carried out after fertilisation (except for E_0_ for which we are not sure whether microdissection took place before or after fertilisation), and hence the male centrioles were present. Nevertheless, we observed cell division and morphological defects when the stalk was removed up to the 8-cell stage. Therefore, MUM is distinct from the male centrioles as well. Altogether, it appears unlikely that MUM corresponds to either centrioles or flagella.

### Is MUM chemical or mechanical ?

#### The stalk is a dead structure composed of an inert cell wall filled with viscous material

The remnant cell wall of the oogonium contains some mucosal compounds (Klochkova et al., 2019, and our own observations). We never observed any intracellular content, such as autofluorescent chloroplasts, or DAPI-stained nuclei, but some authors have reported the presence of some oogonium cytoplasm left-overs in other families of Laminariales (e.g. Alaria: Klimova and Klochkova, 2017). Interestingly, in other families of Laminariales, the basal cell of the embryo (e.g. in *Eisenia bicyclis, Ecklonia* sp.) or the maternal gametophyte itself (in *Undaria* sp.) grows into this empty space once the egg has been released (Kanda, 1941). When the embryo itself grows into the empty stalk, the stalk does not modify its development (Kanda, 1941; Eisenia, plate 37). Therefore, although the stalk appears to be a volume that the embryo can occupy, it is filled with seemingly inert, viscous components. These viscous components are most likely polysaccharides extruded from the stalk cell wall or from the embryo itself. Furthermore, the cell wall of the stalk does not appear to be able to modify cell fate, unlike the cell wall of the rhizoid of the *Fucus* brown alga embryo, as reported by (Berger et al., 1994; Bouget et al., 1998).

#### Is MUM a constricting collar?

The region of the stalk interacting with the E/Z/E coincides with the presence of a thick cell wall collar (Kanda, 1936; Klimova and Klochkova, 2017), which may clasp the basal region of the zygote and subsequently the early embryo, thereby preventing its growth in the Y-axis. However, in control embryos, inhibition of longitudinal division occurs up to the 8-cell stage, when the embryo is 50-60µm long. It is therefore difficult to envisage how a mechanical constraint exerted by a constricting collar at the top of the stalk could act beyond the base of the zygote/embryo itself.

#### Can MUM be polysaccharides accumulated at the stalk-embryo junction?

Instead, the polysaccharides of the stalk cell wall could serve as a source of signalling glycoconjugates. In the kelps *Saccharina* and *Alaria,* a cocktail of polysaccharides, mainly composed of α-D-glucose, α-D-mannose and L-fucose have been shown to accumulate at the junction between the stalk and the egg (Klimova and Klochkova, 2017; Klochkova et al., 2019). This cocktail, which reflects the presence of specific alginates and fucans — being the two main components of the cell wall in brown algae (Charrier et al., 2019) —, can no longer be detected when the egg is free floating, unattached to the stalk or to another substratum. Therefore, the stalk-zygote junction is a specific site where the composition of the cell wall differs from the rest of the surface of the zygote. Could this cocktail be MUM? The role of sugars as signalling molecules involved in development remains to be demonstrated in brown algae, but is well known in other organisms such as plants (Mishra et al., 2022; Wang et al., 2021).

#### MUM diffuses acropetally through Phase I embryos

In intact embryos, the first longitudinal division usually occurs in the upper half of the embryo, suggesting that the first cells out of MUM control are apical cells. This characteristic is observed in many members of Laminariales (Sauvageau, 1918) and of other orders (e.g. *Sacchoriza,* Tilopteridales (Fritsch, 1945; Norton, 1972). This indicates a ubiquitous and acropetal mode of action of MUM, in which the inhibitory effect is strong in the basal part of the embryo, but diminishes towards the apex. In our study, the first longitudinal cell divisions occur at least 40 µm away from the stalk (Fig. 8). Therefore, when the embryo cells are at least 40 µm from the stalk, due to the longitudinal growth of the embryo, they are no longer under the negative control of MUM and begin to divide longitudinally. Hence, MUM’s range would be around 40µm.

A distance-based inhibitory relationship with a signalling molecule is not a foreign concept in brown algae, especially kelps. For instance, the reproductive structures (sori) are formed away from the growing region that surrounds the basal transition zone, due to the action of a sporogenesis inhibitor (Buchholz and Lüning, 1999; Pang and Lüning, 2004), which may be auxin (Kai et al., 2006).

Plasmodesmata, which have been observed in both early (up to the 4-cell stage, not shown) and later stages of *Saccharina* embryogenesis (Theodorou and Charrier, 2023), may contribute to the formation of a symplastically diffusible gradient of molecules originating from the stalk, and forcing transverse cell divisions throughout the embryo. This action could be achieved by any type of compound, as long as its size does not exceed 20–40 kDa (10–20 nm) (Nagasato et al., 2017), which is a much larger size compared with land plant plasmodesmata, but large enough to allow signalling molecules such as auxin to pass through.

Reconsidering all these hypotheses on the nature of MUM, i.e. that of a signal directly (MUM is a glycoconjugate) or indirectly (MUM is a degradation product of glycoconjugate) produced from the viscous content filling the stalk or from its thick collar, the chemical nature of MUM remains the most likely to explain its acropetal diffusion throughout the embryo and its inhibitory effect on longitudinal divisions during 5 days (duration of Phase I) over a distance of around 50 µm. Nevertheless, this hypothesis remains to be confirmed and the exact chemical nature of MUM characterised.

### Evolution of the maternal tissue-embryo connection

The ancestor of the Stramenopiles, the main group the brown algae, diverged from their eukaryote ancestors at least 1 billion years ago (Burki et al., 2020). Among brown algae, at least four orders (Laminariales, Sporochnales, Desmarestiales and to a lesser extent Tilopteridales) display a stalk-mediated, physical connection of the embryo with the maternal tissue that persists over the egg or zygote stages (Fritsch, 1945). Although stalks connecting eggs with maternal tissue are common in Fucales (Burridge et al., 1993), this connection is transient and does not persist after fertilisation.

Compared with other multicellular organisms, this type of interaction is rare in the tree of life. In the green alga *Coleochaete* sp. (Chlorophyta), the zygote remains attached to the female gametophyte until it completes a series of divisions and it releases up to 32 biflagellate spores. This spore release excludes any gametophyte-dependent development of an embryo. Furthermore, this is a case of matotrophy only (Haig, 2015), unlike the case of *Saccharina*.

In several red algae (Gelidiales, Gracilariales), the (carpo)sporophyte develops on the maternal gametophyte, but in the form of gonimoblasts, diploid filaments that produce carpospores, which disperse before developing into diploid (tetra)sporophytes away from the maternal tissue (van der Meer, 1979). Thus, maternal tissue does not directly control sporophyte morphogenesis, and its relationship with the carposporophyte (filamentous gonimoblast) is trophic (matotrophy). The trophic link is particularly evident in Gelidiales and Gracilariales, where nutrient cells and nutrient filaments are found respectively. In the red alga *Palmaria palmata*, however, the situation is similar to that of *Saccharina*: the embryo develops as a macroscopic elongated blade, in physical contact with dwarf haploid maternal tissue (Gall et al., 2004; van derMeer and Todd, 1980). However, it is not known whether the latter has an impact on the development of the embryo’s growth axes.

In contrast, in bryophytes (group comprising the mosses, liverworts and hornworts), the developing embryo is surrounded by a layer of maternal cells, making up the archegonium (Naf, 1962) and subsequently covered by the calyptra, a cell layer of female origin protecting the embryo from desiccation (Budke et al., 2012). First, the embryo is formed and develops nested within the archegonium cavity, and then emerges out of it. Subsequent evolution of this branch led to increasingly protected embryos surrounded by additional layers of maternal cells, as in the seeds of land plants, where the embryo is embedded in albumen, a triploid tissue resulting from the fertilisation of a diploid maternal tissue, and the integuments and fruit differentiating from maternal tissues. Although phylogenetically very distant, this latter phanerogam embryo development is reminiscent of the viviparous mode of embryo production in animals, where the embryo remains nested in the maternal body up to its mature stage. In contrast, other organisms, including all other brown algae, grow their embryo outside the parental tissues. In *Fucus* (order Fucales), female and male gametophytes release eggs and sperm cells into the seawater where fertilisation takes place (Hatchett et al., 2022), as in fish, amphibians and many aquatic invertebrates such as tunicates and echinoderms (ovipary). Therefore, *Saccharina* represents an intermediate case, and one can reasonably hypothesise that, in view of the fragile physical connection between its egg and the maternal stalk (eggs detach from the stalk even with gentle shaking), this mode of interaction will evolve towards either vivipary-like or ovipary-like embryo development. That Laminariales, Tilopteridales, Sporochnales and especially the Desmarestiales, which diverged early (Bringloe et al., 2020; Silberfeld et al., 2010), are pioneers of a parental-embryo physical connection in brown algae distinct from ovipary is congruent with the fact that they diverged independently during the radiation giving rise to most current brown algal orders. In any case, this particular mode of maternal-embryo interaction has clearly enabled Laminariales to become the largest and most morphologically complex brown algae to thrive in the oceans.

## Materials and Methods

### Algal culture

The culture and production of embryos of *Saccharina latissima* (Arthrothamnaceae, Laminariales, Phaeophyceae) were carried out according to Theodorou et al. (2021). Female (F1) and male (M1) gametophytes with a fixed genotype were used to produce all the embryos. These genotypes were selected from the offspring of one mature sporophyte collected on the beach at Perharidy (Roscoff, Brittany, France) (48°43’33.5"N 4°00’16.7"W) based on their growth rate and sexual compatibility when cultured *in vitro*. F1 and M1 gametophytes were amplified by vegetative multiplication as described in Theodorou et al. (2021). Gametes were obtained from the maturation of gametophytes under 16 μmol photons m^-2^·s^-1^ white light intensity and 14:10 light:dark photoperiod at 13 °C. Embryos were observed after transferring the cultures to higher light intensity (50 μmol photons m^-2^·s^-1^) for a week.

### Excision of the maternal tissue at different developmental stages

The separation of embryos from their stalk was carried out by using pulled glass micro-needles. First, glass micro-needles were prepared by pulling glass capillaries (GC100F-10) with a pipette puller (SU-P97 Flaming/Brown type micropipette puller) using the following programme: heat, 564°C; pull, 70 U; velocity, 70 ms; time, 250 ms. After pulling, the tip of the needle was sharpened to ensure precision cutting. Second, developing embryos were selected under a flow hood using an inverted microscope (Olympus CKX41 Inverted with phase contrast). Then, the tip of the stalk was cut using the glass needle in a cutting motion, while holding the embryo down on the bottom of the dish. This action was repeated for eggs, zygotes, 2-cell, 4-cell and 8-cell embryos, as well as early Phase II embryos (corresponding to 8-cell to ∼ 1000 cell embryos, see Theodorou & Charrier, 2023 for the definition of the embryogenesis stages). For use as controls, i) whole gametophytes, ii) gametophytes with damaged cells distant from the stalk, and iii) embryos attached to the substratum (bottom of the plastic Petri dish), were transferred to another dish using a non-pulled capillary and a manual microinjector (Eppendorf CellTram Air 5176).

### Cell staining

#### Trypan blue

A drop of Trypan blue was added to fertile female gametophytes of *S. latissima* immersed in ∼ 0.5 mL of seawater, followed by observation in bright field microscopy (DMI8, Leica Microsystems).

#### Calcofluor white

Staining of cell walls from embryos separated from the maternal tissue was performed at different times post excision. Embryos were fixed for 1 h in equal parts of 4% PFA in H_2_O and filtered natural seawater (NSW). After fixation, the samples were washed in NSW and twice in PBS to remove any excess fixative. Subsequently, they were incubated with 20 µM Calcofluor white (Sigma-Aldrich) for 3 days at 4°C in the dark. Following incubation, the samples were washed three times with PBS to remove any unbound dye. Finally, the algal samples were mounted using Cityfluor mounting medium (Electron microscopy sciences).

### Image acquisition

All experiments of time-lapse microscopy of growing embryos were recorded under a bright-field microscope (Leica DMI600 B, Olympus CKX41 or Leica DMi8 inverted phase contrast microscopes) equipped with a DFC450C camera with acquisition intervals of 2 to 24 h for a duration of at least 10 days after excision. The required temperature and light were set in a carbonate-glass chamber fitted to the microscope and equipped with a thermostatically controlled airflow system (Cube and Box, Life Imaging services) and commercially available LED white light sources. Observations of cell wall staining (see experimental procedure for Calcofluor white staining below) were performed with a Leica TCS SP5 AOBS inverted confocal microscope (20X objective / N.A. 0.70 and correction 0; Exc/Em band wavelengths: 405/561-596 nm; pinhole, 60.6 µm).

### Manual segmentation

Automatic segmentation proved to be challenging due to the constant pigmentation changes of the cells transitioning from high to low coloration. Additionally, daily exposure to UV light required to display Calcofluor white staining proved to be detrimental to the algae. Therefore, manual segmentation was carried out from bright-field images. To minimise image deformation, flat-growing embryos were predominantly chosen. Z-stack images of time-lapse acquisition were segmented manually with Fiji (ImageJ2 version 2.9.0) (Schindelin et al., 2012) and the outlines were implemented in Inkscape (version 1.2). Resulting cell wall contours were analysed using in-house software (see below). The programme extracted multiple quantitative parameters for each embryo lamina (blade) and its cells.

### Quantitative morphometry

Cell wall vector graphics were processed in dedicated software written in object-oriented python 3 (Van Rossum and Drake, 2023) that we called blade_painter. Reading the svg file, blade_painter extracts various geometric properties for cells and laminae (Rosin, 2005), namely (a) each cell contour, from which were directly derived the perimeter length and surface area; (b) convex hull, used to compute the minimal bounding rectangle (MBR). The main axis and length/width (l/w) ratio (or elongation) of the cell were assumed to be those of the MBR; (c) rectangularity, computed as the proportion of overlapping surface between the cell and its MBR, rescaled to the same area; (d) neighbour cell count, where two cells were considered neighbours if they shared at least 200 nm of cell wall (this threshold can be modified as a software parameter); (e) blade area; (f) main and secondary axes of orientation and length of the blade, computed according to Fletcher et al. (2013).

### Statistical analysis

Data collected from the segmented cell and blade contours were analysed using standard python3 libraries, namely pandas (The pandas development team, 2020) (pandas-dev/pandas: Pandas) and scipy (Virtanen et al., 2020). All pairwise comparisons for cell data were conducted using the Student’s mean comparison test with Welch’s correction, except for neighbour cell numbers and orientation, which were compared using a χ² test. Blade data were compared using the nonparametric Mann-Whitney test.

### Growth rate

For each observed blade, the date of observation was set to *t*=0 at the transition from Phase I to Phase II (first longitudinal division). Linear regression was performed on the logarithm of cell number as a function of time, for - 48 ≤ *t* ≤ 72. The increase rate was computed as *r* in the equation *N*=*N*_0_ ×*r^t^*, thus derived from the slope of the regression line log (*N*)=log (*N*_0_)+log (*r*)×*t*. From *r*, we inferred the doubling time *τ* =1/ log_2_(*r*).

## Acknowledgements

S.B was funded by the ARED PhD programme from Bretagne Regional Council (“Primaxis” project, grant number 1749) and Sorbonne University. I.T was funded by the ARED PhD programme from the Bretagne Regional Council (“PUZZLE” project) and NMBU. Part of the work was funded by the European Union (ERC, ALTER e-GROW, project number 101055148). Views and opinions expressed are however those of the author(s) only and do not necessarily reflect those of the European Union or the European Research Council Executive Agency. Neither the European Union nor the granting authority can be held responsible for them. For purposes of open access dissemination, a CC-BY-NC license has been applied by the authors on this document. We thank Bruno de Reviers for fruitful discussion about reproduction of brown algae.

## Supplementary figures

**Suppl. Fig. 1.**
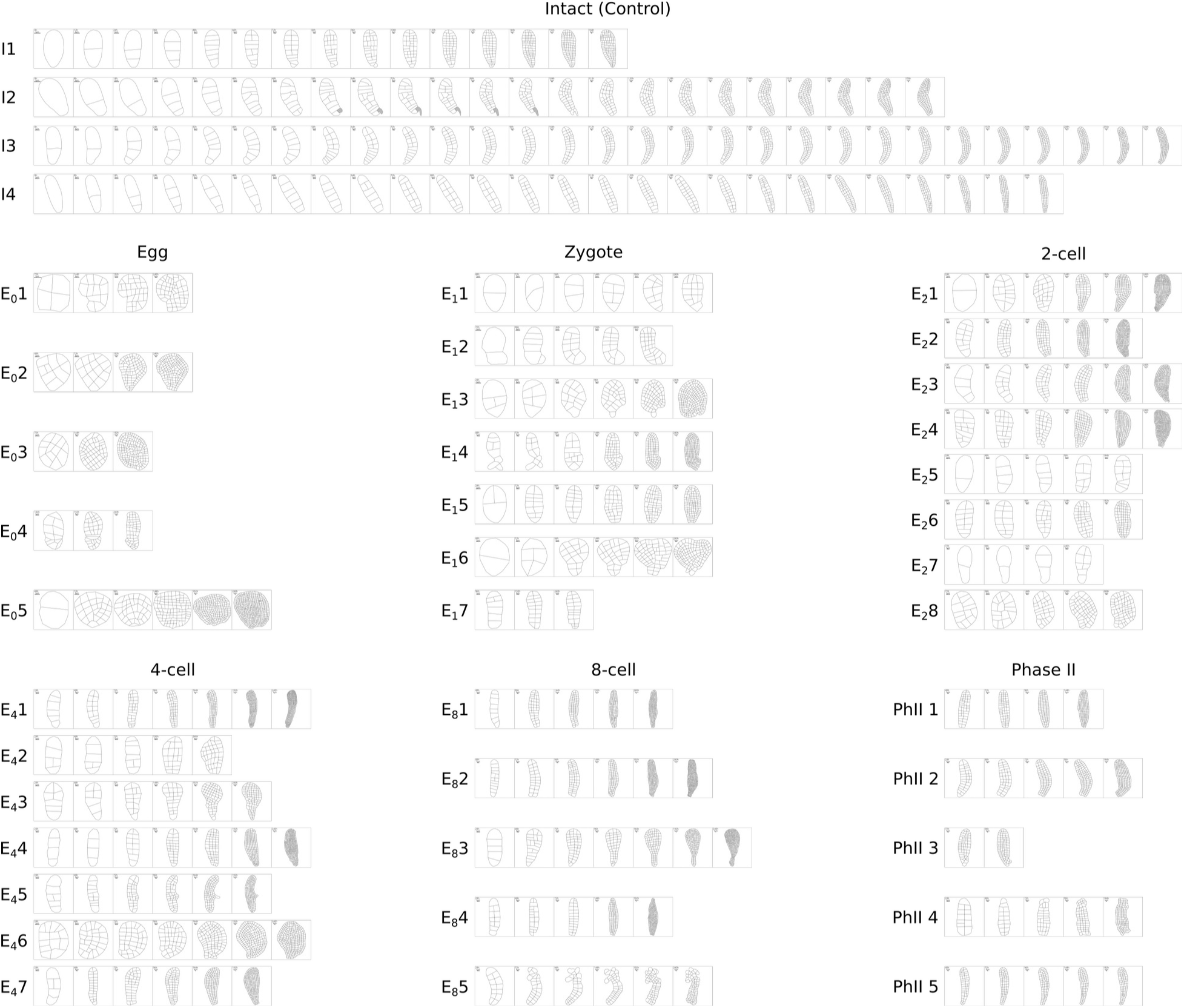
Results of manual segmentation of the embryos. The cell outlines were drawn manually from a series of bright-field z-stack images (Stack focuser, see Materials and Methods section) and used for quantitative analyses of the morphometry parameters. The stages at which the separation from the stalk took place is indicated with the code used in the article (e.g. E_0_ for the separation from the stalk at the egg stage). I is for intact embryos. Replicates are indicated with number.

**Suppl. Fig. 2.**
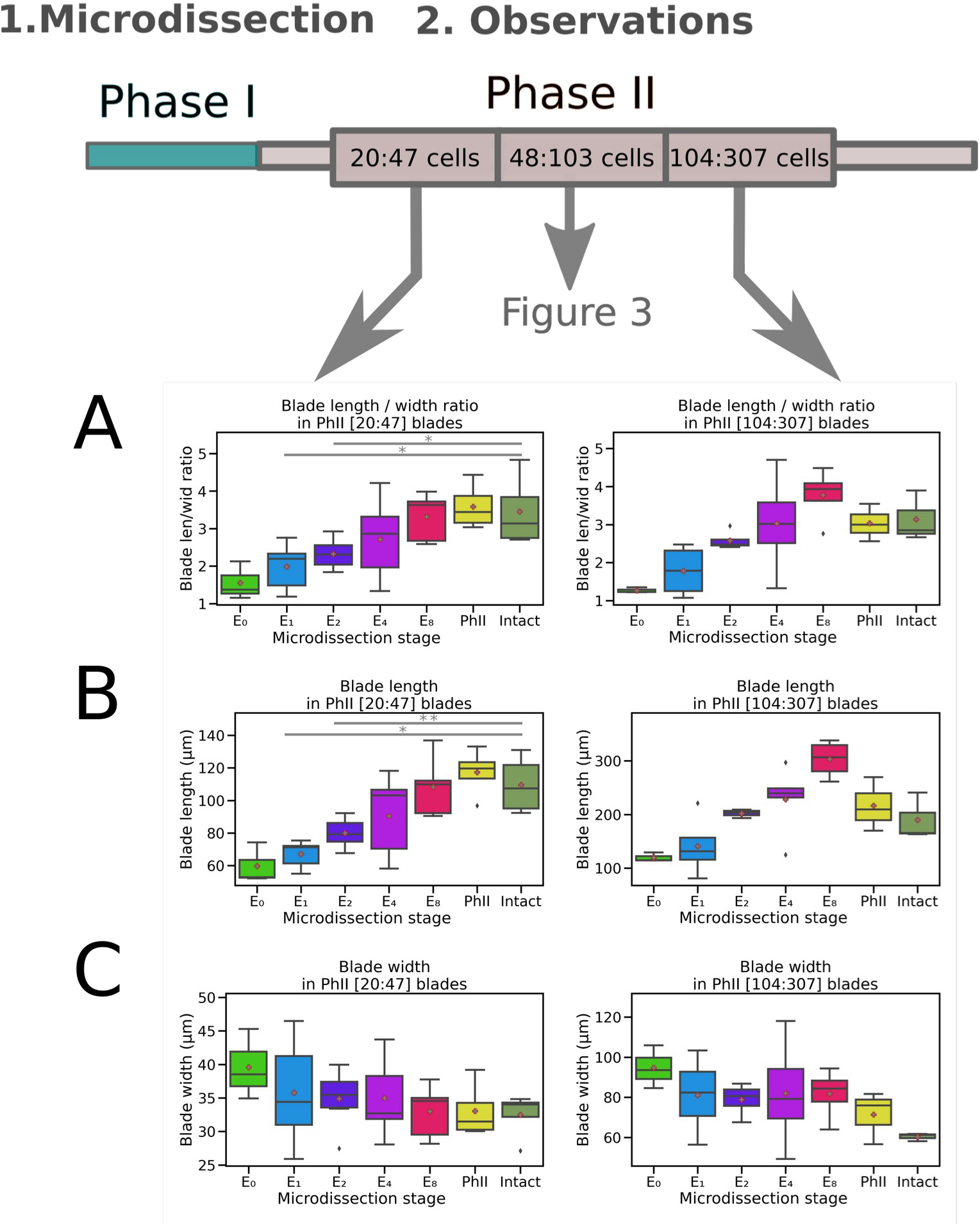
Impact of embryo detachment from the stalk on blade shape at two additional developmental windows. The distribution of the length/width ratios of blade of different stages (20–47-cell, left and 104– 307-cell embryos, respectively left and right panels) grown after microdissection of the stalk at the E_0_, E_1_, E_2,_ E_4_, E_8_ and PhII stages was plotted and compared with intact embryos. The line in the box shows the median, the box outline frames the first quartile and the whiskers indicate the third quartile. Results of the *t*-test are indicated by p-values: *<5.10^-2^; ** <1.10^-2^; *** < 10^-3^. This figure supplements Fig. 3.

**Suppl. Fig. 3.**
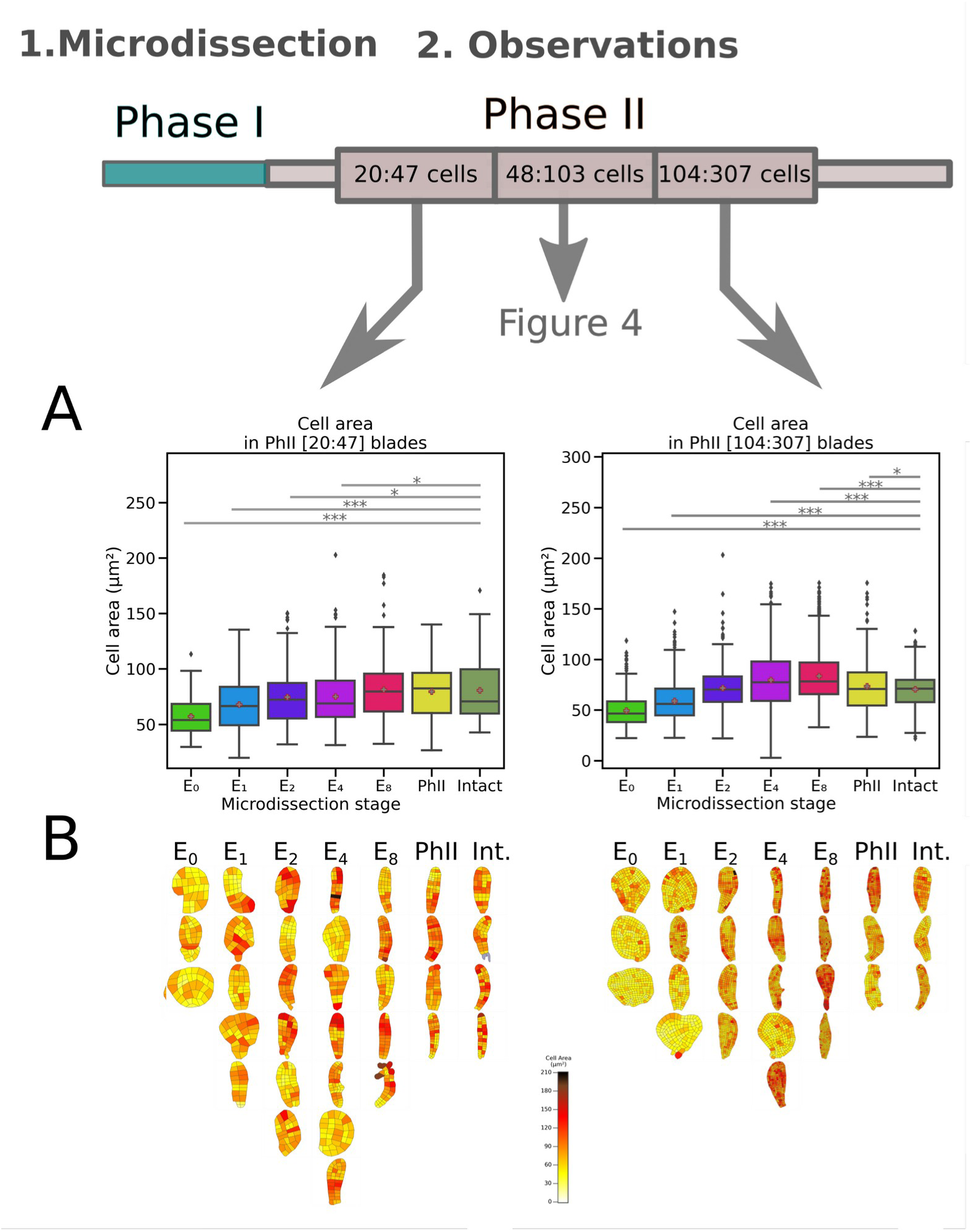
Impact of detachment from the stalk on cell areas at two additional developmental windows. (A) The distribution of the cell areas of embryos at different developmental stages (20–47-cell, and 104–307-cell embryos, respectively left and right panels) grown after microdissection of the stalk at E_0_, E_1_, E_2,_ E_4_, E_8_ and PhII stages was plotted and compared with intact embryos. The line in the box shows the median and the box outline frames the first quartile. The asterisk indicates the mean. The whiskers indicate the third quartile. Results of the *t*-test are indicated by p-values: *<5.10^-2^; ** <1.10^-2^; *** < 10^-3^. This figure supplements Fig. 4B. (B) Heatmap of cell areas for each embryo considered in (A). Colour scale indicated on the bottom right-hand side of the figure is specific to the corresponding time window and may be different from that shown in Fig. 4. The scale is adjusted so that the image of each embryo (blade) has the same dimensions.

**Suppl. Fig. 4.**
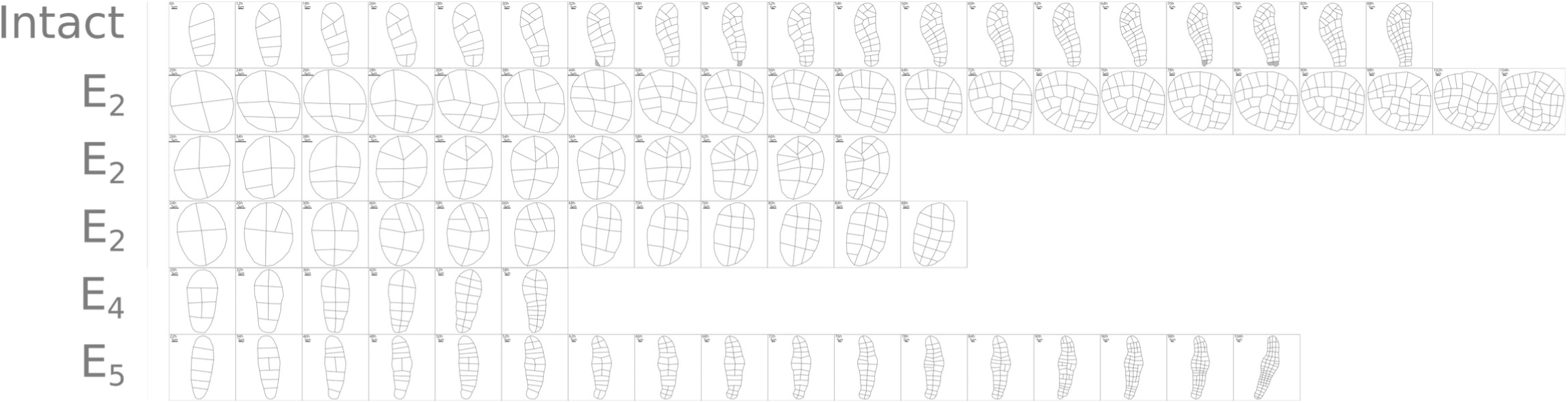
Time series of microdissected embryos. Photos of embryos imaged every 2 h were segmented at each additional cell division (the time interval between cell division is about 24 h, so that not all the images were segmented). Each line represents one embryo, the stage at which microdissection took place is indicated in the left margin. E_5_ is a 5-cell stage embryo. Scale bar is indicated on each embryo. Note that it differs mainly between intact and microdissected embryos.

**Suppl. Fig. 5.**
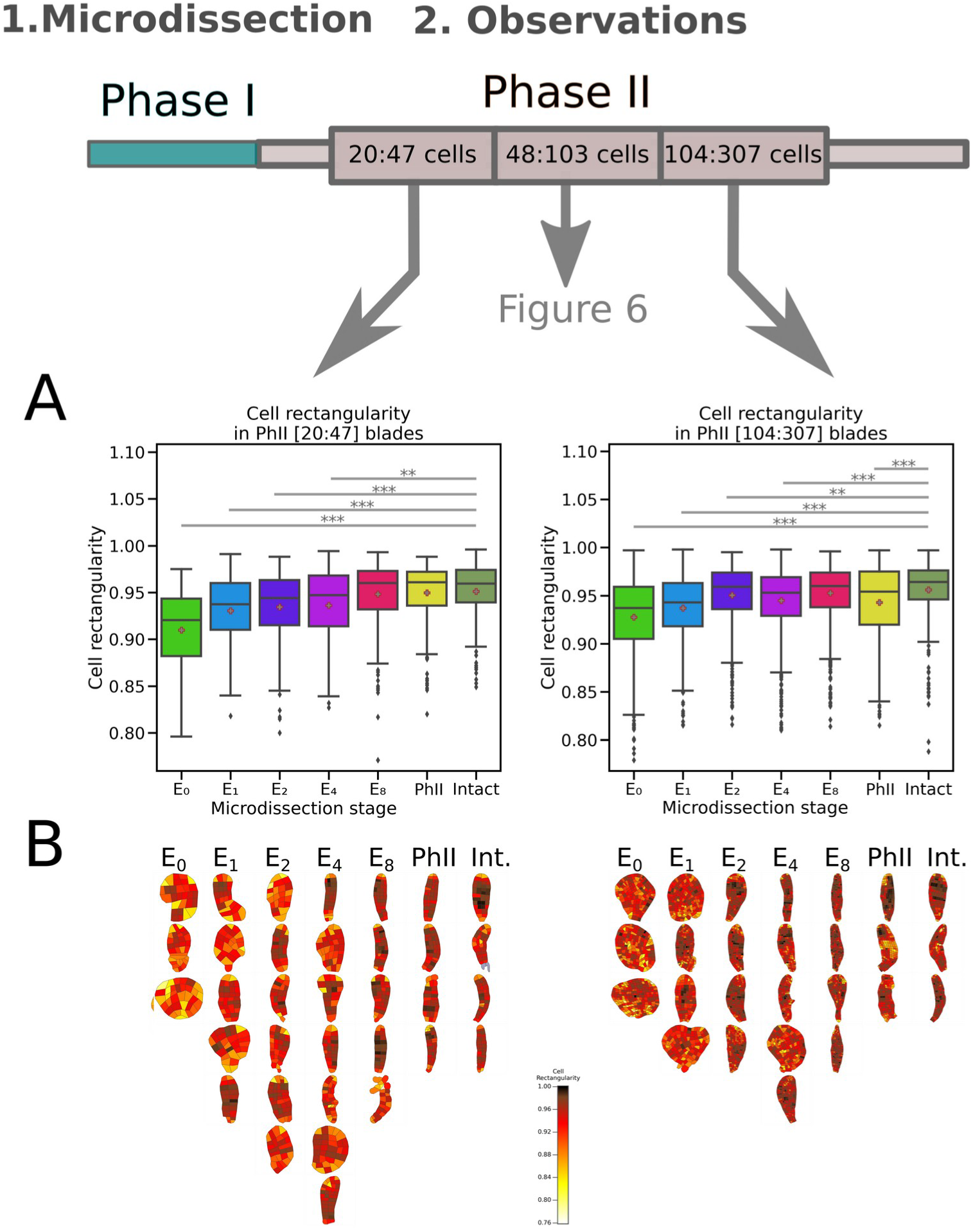
Impact of detachment from the stalk on embryo cell shapes at two additional developmental windows. Cell shape is expressed as a factor of rectangularity. (A) The distribution of the cell rectangularity (see Methods for definition) of embryos of different stages (20–47-cell and 104– 307-cell embryos, respectively left and right panels) grown after microdissection of the stalk at E_0_, E_1_, E2, E_4_, E_8_ or PhII stages was plotted and compared with intact embryos. The line in the box shows the median, the box outline frames the first quartile and the whiskers indicate the third quartile. Asterisks show the mean. Results of the *t*-test are indicated by p-values: *<5.10^-2^; ** <1.10^-2^; *** < 10^-3^. This figure supplements Fig. 6. B. (B) Heatmap of cell rectangularity for each embryo considered in (A). Colour scale indicated on the bottom right-hand side of the figure is specific to the corresponding time window and may be different from that shown in Fig. 6. The scale is adjusted so that the image of each embryo (blade) has the same dimensions.

**Suppl. Fig. 6.**
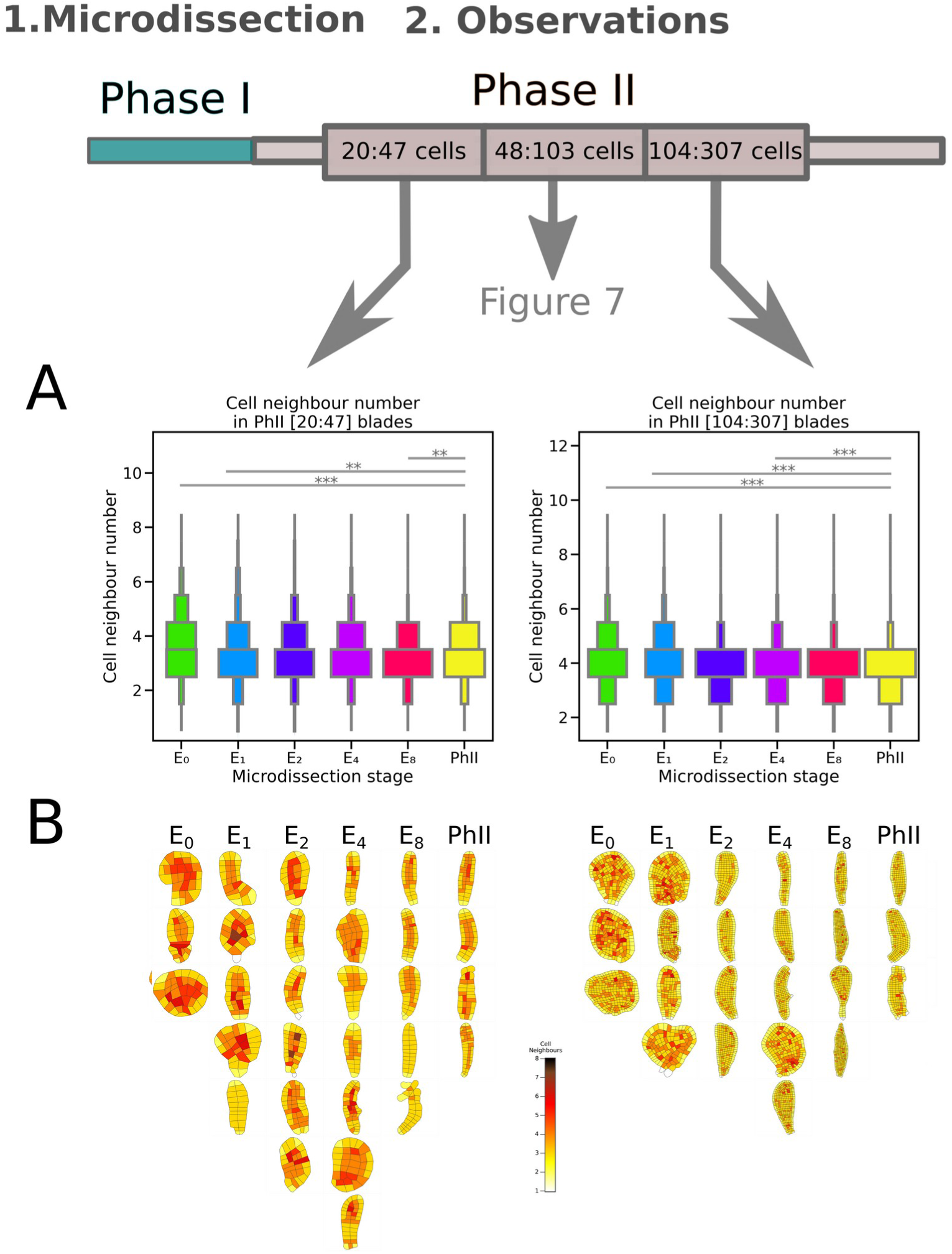
Impact of dissection from the stalk on the number of cell neighbours in microdissected embryos when observed at two different developmental windows. (A) The distribution of the number of cell neighbours in embryos of different stages (20–47-cell and 104–307-cell embryos, respectively left and right panels) grown after microdissection of the stalk at E_0_, E_1_, E2, E_4_, E_8_ or PhII stage was plotted as violin plots and compared with intact embryos. Results of the Chi^2^-test are indicated by p-values: *<5.10^-2^; ** <1.10^-2^; *** < 10^-3^. This figure supplements Fig. 7. (B) Heatmap of cell areas for each embryo considered in (A). Colour scale indicated on the bottom right-hand side of the figure is specific to the corresponding time window and may be different from that shown in Fig. 7. The scale is adjusted so that the image of each blade has the same dimensions.

## Additional material

**Suppl. Table 1**. Morphometric parameters of the embryo (blade). The table is sub-divided in 3 sections based on the size of embryos (expressed in the number of cells nCells, first and 4th columns).“In" are intact embryos. Not all embryos have available data for all three size ranges.

**Suppl. Table 2.** P-values of statistical tests comparing morphometric parameters characterising *Saccharina* embryos and their cells. *t*-test were used to compare the cell features, Wilcoxon tests to compare embryo blades and Chi² for the analyses of the number of cell neighbours. The quantitative value of each parameter and for each developmental stage at which the stalk was sectioned (En and PhII), is presented as the function of the time window (expressed as the number of cells). Only p-values resulting from the comparison between the microdissected embryos and the intact control are shown in the main figures presented in the article. In green, p-Values lower than 5.10^-2^.

**Suppl. Table 3.** Cell morphometrics in each embryo (blade).Time point and number of cells are indicated for each embryo. Each cell (one row) was studied for different morphometric parameters. Those relevant for this study are displayed. The data are shown for the three ranges of embryo size (in cell number; e.g. [20:47] cells, indicated by the first column).

**Suppl. Movie 1.** Development when egg naturally detached from the stalk

